# The *ROP16_III_*-dependent early immune response determines the sub-acute CNS immune response and type III *Toxoplasma gondii* survival

**DOI:** 10.1101/645390

**Authors:** Shraddha Tuladhar, Joshua A. Kochanowsky, Apoorva Bhaskara, Yarah Ghotmi, Anita A. Koshy

## Abstract

*Toxoplasma gondii* is an intracellular parasite that persistently infects the CNS and that has genetically distinct strains which provoke different acute immune responses. How differences in the acute immune response affect the CNS immune response is unknown. To address this question, we used two persistent *Toxoplasma* strains (type II and type III) and examined the CNS immune response at 21 days post infection (dpi). Contrary to acute infection studies, type III-infected mice had higher numbers of total CNS T cells and macrophages/microglia but fewer alternatively activated macrophages (M2s) and regulatory T cells (Tregs) than type II-infected mice. By profiling splenocytes at 5, 10 and 21 dpi, we determined that at 5 dpi type III-infected mice had more M2s while type II-infected mice had more classically activated macrophages (M1s) and these responses flipped over time. To test how these early differences influence the CNS immune response, we engineered the type III strain to lack ROP16 (IIIΔ*rop16*), the polymorphic effector protein that drives the type III-associated M2 response. IIIΔ*rop16*-infected mice showed a type II-like neuroinflammatory response with fewer infiltrating T cells and macrophages/microglia and more M2s and an unexpectedly low CNS parasite burden. At 5 dpi, IIIΔ*rop16*-infected mice showed a mixed inflammatory response with more M1s, M2s, T effector cells, and Tregs, and decreased rates of infection of peritoneal exudative cells (PECs). These data suggested that type III parasites need the early ROP16-associated M2 response to avoid clearance, possibly by the Immunity-Related GTPases (IRGs), IFN-γ dependent proteins essential for murine defenses against *Toxoplasma*. To test this possibility, we infected IRG-deficient mice and found that IIIΔ*rop16* parasites now maintained parental levels of PECs infection. Collectively, these studies suggest that, for the type III strain, *rop16_III_* plays a key role in parasite persistence and influences the sub-acute CNS immune response.

**Author Summary:** *Toxoplasma* is a ubiquitous intracellular parasite that establishes an asymptomatic brain infection in immunocompetent individuals. However, in the immunocompromised and the developing fetus, *Toxoplasma* can cause problems ranging from fever to chorioretinitis to severe toxoplasmic encephalitis. Emerging evidence suggests that the genotype of the infecting *Toxoplasma* strain may influence these outcomes, possibly through the secretion of *Toxoplasma* strain-specific polymorphic effector proteins that trigger different host cell signaling pathways. While such strain-specific modulation of host cell signaling has been shown to affect acute immune responses, it is unclear how these differences influence the sub-acute or chronic responses in the CNS, the major organ affected in symptomatic disease. This study shows that genetically distinct strains of *Toxoplasma* provoke strain-specific CNS immune responses and that, for one strain (type III), the acute and sub-acute immune responses and parasite survival are heavily influenced by a polymorphic parasite gene (*rop16_III_*).

## Introduction

*Toxoplasma gondii* is a ubiquitous obligate intracellular parasite that chronically infects the brain, heart, and skeletal muscle of humans (1,2). Up to one third of the world’s population is estimated to be chronically infected with *Toxoplasma* (3). While most infected people are asymptomatic, in some immunocompromised individuals and developing fetuses *Toxoplasma* can cause fever, chorioretinitis, toxoplasmic encephalitis, and even death (4). While the host immune status plays a key role in determining disease outcomes, clinical data suggest that the genotype of the infecting *Toxoplasma* strain may also play a role (5–14). *Toxoplasma* strains are classified into 15 genetic haplotypes which include the three canonical strains — type I, type II, and type III (now haplotype 1, 2, and 3 and Clade A, D, and C respectively) (15–17). Of the canonical strains, type I and III are relatively genetically similar compared to type II.

Our understanding of how different *Toxoplasma* strains might cause distinct disease outcomes in mice and potentially humans has greatly expanded in the last decade. We now know that *Toxoplasma* highly manipulates host cells through the injections and secretion of effector proteins that can be polymorphic among strains. In turn these polymorphisms can profoundly affect the host cell response. For example, during acute *in vitro* infection of fibroblasts or immune cells only the type I/III allele of ROP16 (ROP16_I/III_), not the type II allele, causes direct and prolonged phosphorylation of the transcription factors STAT3 and STAT6 (18,19). In macrophages, this prolonged activation of STAT3/6 leads to decreased production of IL-12, a key pro-inflammatory cytokine (19). Conversely, only the type II allele of GRA15 (GRA15_II_), not the type I/III allele, activates the transcription factor, NFκB, which leads to host cell production of pro-inflammatory cytokines (20,21). In addition, strains that express Gra15_II_ polarize infected macrophages to a classically activated phenotype whereas, strains that express ROP16_I/III_ polarize macrophages to an alternatively activated phenotype (19,21,22). Yet, how these strain-specific modulations of infected cells influence global immune responses, or sub-acute or chronic immune responses in the central nervous system (CNS), remains unknown. The only studies looking at *Toxoplasma* strain-specific tissue immune responses during chronic infection were done 20 years ago and used histology to define the CNS immune response. While these studies identified strain-specific neuroinflammatory responses, the strains also produced different CNS parasite burdens making it impossible to determine if the immune response differences were simply driven by the differences in parasite burden (23,24).

To address this gap and leverage our increased understanding of strain-specific effects, we infected mice with a representative strain from either of the two canonical, encysting *Toxoplasma* strains (type II or type III) and then defined the neuroinflammatory response using quantitative immunohistochemistry (IHC), multiplex cytokine analysis, and flow cytometry. At 21 days post infection (dpi), compared to type II-infected mice, type III-infected mice showed a higher number of macrophages, infiltrating T cells, and levels of pro-inflammatory cytokines (e.g IFN-γ) in the CNS, even though type II and type III-infected mice showed the same CNS parasite burden. In addition, our flow cytometry analyses of CNS and splenic mononuclear cells showed that type III-infected mice had fewer alternatively activated macrophages (M2s) and regulatory T cells (Tregs) compared to type II-infected mice, the opposite of what is seen with acute infection *in vivo* and *in vitro* (21,22). By examining the peripheral macrophage immune response over time, we determined that, early in infection, type III-infected mice have more M2s compared to type II-infected mice and that this response changes over time, leading to fewer M2s in the spleen and brain of type III-infected mice by 21 dpi. To define if the differences in the early macrophage response influenced the subsequent CNS immune response, we engineered the type III strain to lack the *rop16* gene (IIIΔ*rop16*), which, as noted above, is the driver of the early type III-associated M2 response (19,21). Consistent with our hypothesis, compared to the parental type III strain, IIIΔ*rop16*-infected mice showed a more type II-like CNS immune response with fewer macrophages and infiltrating T cells and an increase in M2s in the CNS. Unexpectedly, the IIIΔ*rop16*-infected mice showed a substantial decrease in the CNS parasite burden; a mixed acute inflammatory immune response (i.e. an increase in classically activated macrophages (M1s), M2s, T effector cells, and Tregs); and rapid clearance from the site of inoculation. As the type III strain is sensitive to destruction by Immunity-Related GTPases (IRGs) (25,26), these results suggested that, to persist, the type III strain requires *rop16_III_* to dampen the initial immune response, including the IRG response. We tested this possibility by infecting mice that lack the IRG response (27) and found that the IIIΔ*rop16* strain now maintained parental levels of infection at the site of inoculation. Collectively these data suggest that *Toxoplasma* strain-specific immune responses persist in the sub-acute phase of disease and that, for the type III strain, *rop16_III_* is required for persistence and plays a role in determining acute and sub-acute systemic and CNS immune responses.

## Results

### Type III-infected mice have an increased CNS T cell and macrophage/microglia response compared to type II-infected mice

To determine if genetically divergent *Toxoplasma* strains cause strain-specific CNS immune responses, we infected mice with either a type II (Prugniaud) or type III (CEP) strain and analyzed the CNS macrophage and T cell immune response at 21 dpi, which we consider a sub-acute time point of CNS infection. We focused on the macrophages/microglial and T cell responses because prior work has established that these cells are essential for controlling acute and chronic toxoplasmosis (1,28–31). To quantify macrophages/microglia and T cells in the CNS after *Toxoplasma* infection, we stained brain sections with antibodies against Iba-1, a pan macrophages/microglial marker, or CD3, a pan-T cell surface marker. Stained sections were then analyzed by light microscopy. We found that type III-infected mice had approximately twice the number of CNS macrophages/microglia compared to type II-infected mice (Fig. 1A,B). Type III-infected mice also had a similar increase in the number of CNS T cells (Fig. 1C,D).

**Fig 1.**
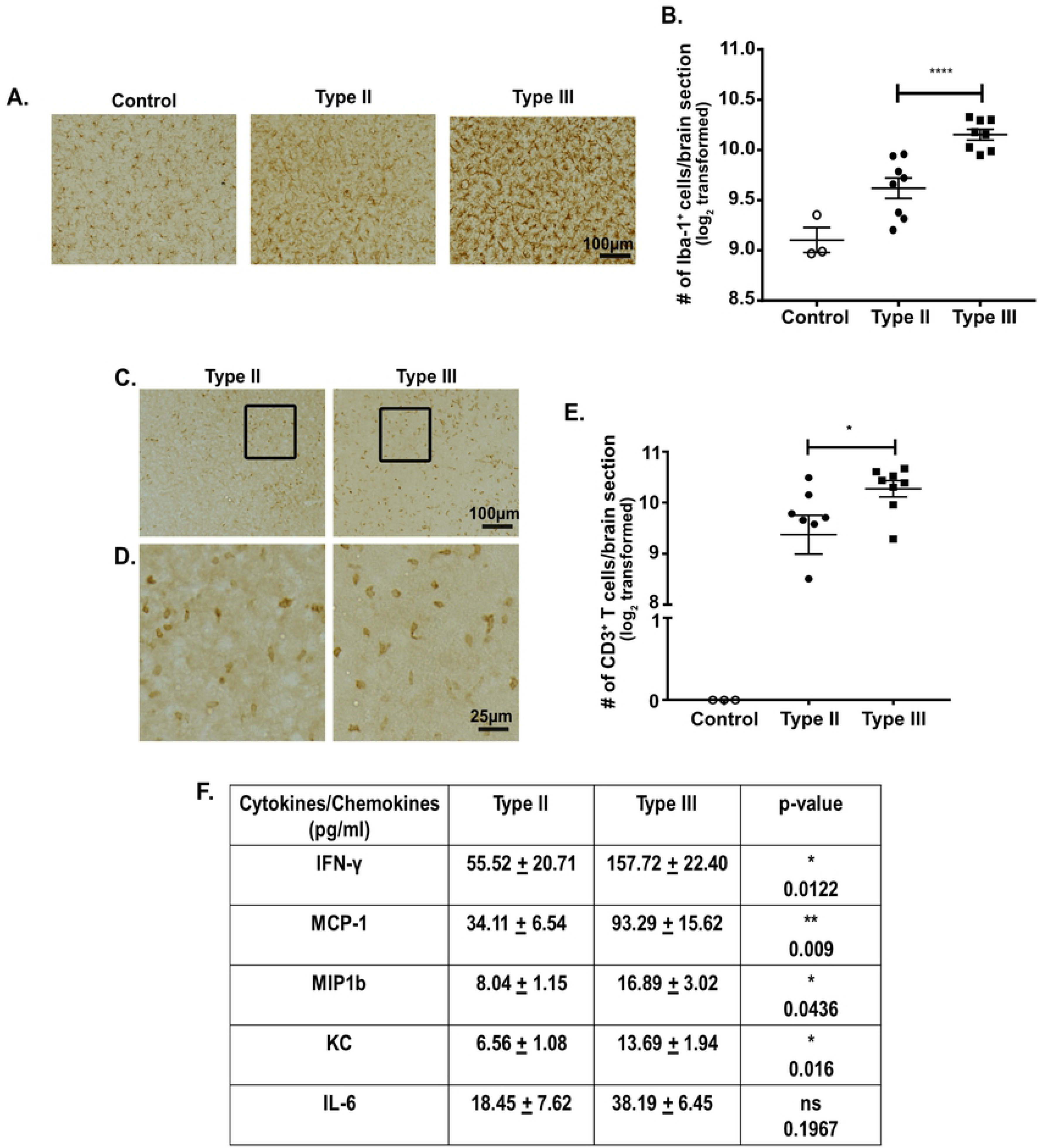
Type III infection provokes a stronger pro-inflammatory CNS immune response compared to type II infection. Mice were inoculated with saline (control), type II, or type III parasites and brains were harvested at 21 days post infection (dpi). **A.** Representative images of Iba-1^+^ cells (macrophages/microglia) in brain sections from mice in each group. **B.** Quantification of the number of Iba-1^+^ cells. **C.** Representative images of CD3^+^ cells (T cells) in brain sections from type II, and type III-infected mice. **D.** Enlargement of the boxed area in **C**. **E.** Quantification of the number of CD3^+^ cells. For **B,E.** Bars, mean ± SEM. N=12 fields of view/section, 3 sections/mouse, 8 mice/infected group. For each mouse, the number of cells/section was averaged to create a single point. *p<0.05, ****p<0.0001, Two-way ANOVA with Fisher’s protected LSD. Data are representative of 3 individual experiments. **F.** The table lists the subset of cytokines or chemokines from a 25-plex assay that showed a ≥2-fold difference between protein levels in brain homogenates from type II and type III-infected mice. Table S1 has full list of cytokines/chemokines. N=8 mice/infected group. p-values determined by one-way ANOVA with Bonferroni post-hoc test.

To determine how the influx of these immune cells changed the global CNS cytokine/chemokine environment, we isolated and analyzed protein from brain homogenates of control (saline inoculated) or infected mice using a 25-plex cytokine and chemokine LUMINEX assay. As expected, compared to control mice, type II and type III-infected mice showed a ≥2-fold increase in most of the pro-inflammatory cytokines and chemokines in the panel (**Table S1**). A subset of these cytokines and chemokines also showed a ≥2-fold increase in type III-infected mice compared to type II-infected mice (Fig. 1F).

Together, these data show that, at 21 dpi, type III-infected mice have significantly higher numbers of both macrophages/microglia and T cells in the CNS as compared to type II-infected mice. Consistent with this increase in CNS immune cells, type III-infected mice have a stronger CNS pro-inflammatory cytokine and chemokine milieu compared to type II-infected mice.

### Differences in the CNS immune response between type II and type III-infected mice are not driven by parasite burden

Given the consistent differences we found in the number of macrophages/microglia and T cells (Fig. 1), we sought to determine if these differences simply reflected disparities in parasite burden. To address this question, we analyzed CNS parasite burden by two methods. First, using DNA isolated from brain homogenates, we performed quantitative PCR (qPCR) for the *Toxoplasma*-specific gene B1 (32–35). Second, we quantified CNS cyst numbers by performing immunofluorescent assays on brain sections using Dolichos biflorous agglutinin (DBA), a lectin that stains the cyst wall, to identify cysts (36). By both measures, we found that type II and type III-infected mice had equivalent CNS parasite burdens at 21 dpi (Fig. 2A,B).

**Fig 2.**
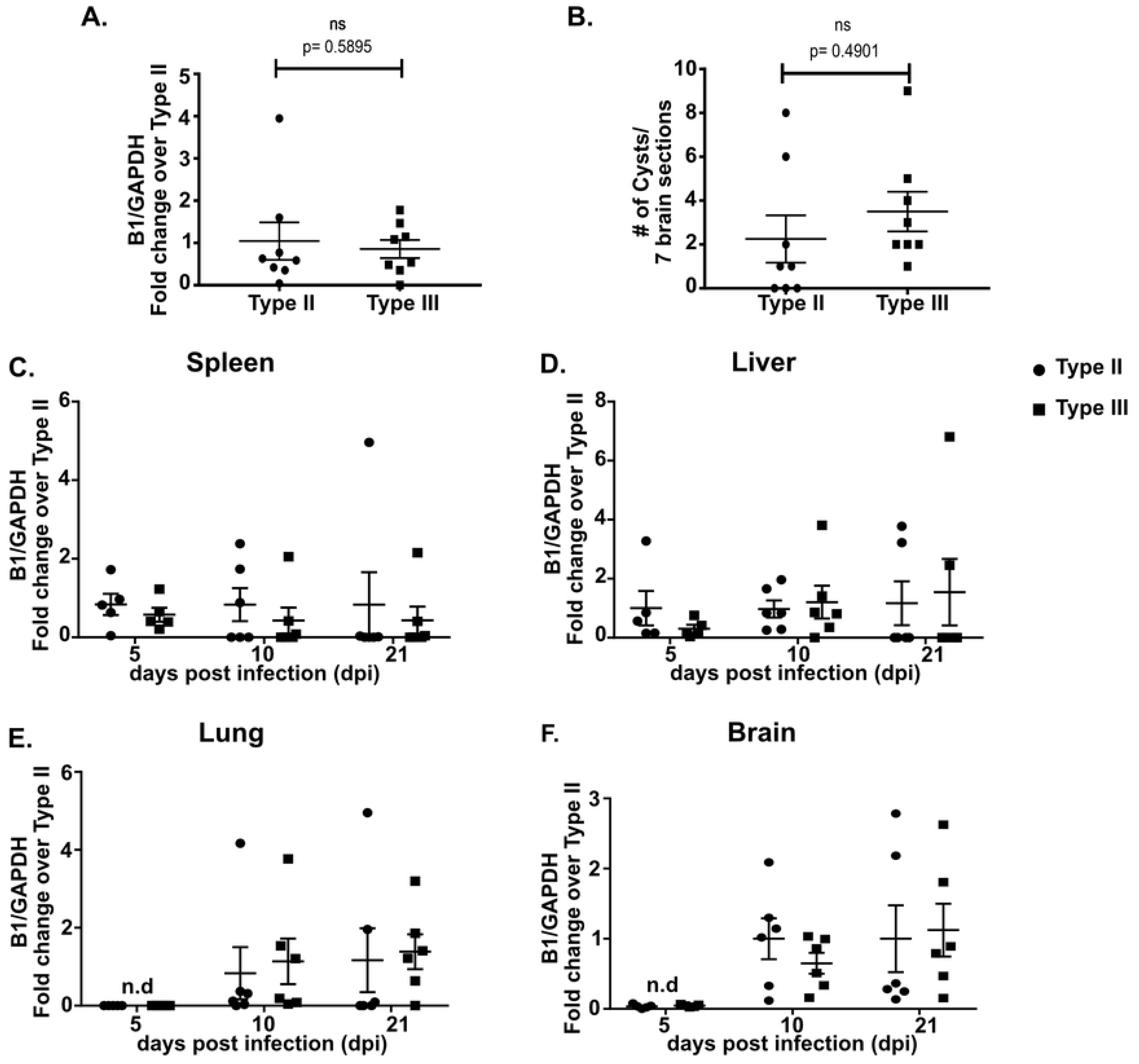
Type II and type III-infected mice show similar dissemination pattern and parasite burdens in early and sub-acute CNS infection. Mice were inoculated with type II or type III parasites, and brains were harvested at noted time points. **A.** Quantification of CNS *Toxoplasma* burden at 21 dpi using quantitative PCR (qPCR) for the *Toxoplasma*-specific B1 gene and host GAPDH gene (housekeeping gene). *Toxoplasma* and mouse genomic DNA were isolated from brain homogenates. **B.** Quantification of *Toxoplasma* cyst burden at 21 dpi in brain sections stained with Dolichos biflorous agglutinin (DBA), which stains the cyst wall. Stained sections were then analyzed by epifluorescent microscopy to quantify DBA^+^ mCherry^+^ cysts. For **A,B**. Bars, mean ± SEM. N= 8 mice/infected group. ns= not significant; two-way ANOVA with Fisher’s protected LSD. Data representative of 3 individual experiments. **C,D,E,F.** Quantification of *Toxoplasma* burden was performed as in (**A**) at specified time points from spleen, liver, lung, and brain. Bars, mean ± SEM. N=4-5 mice/infected group/time point. No significant differences were found in mean B1 quantification. Two-way ANOVA with Fisher’s protected LSD. Data are representative of 2 individual experiments. • = type II, ■ = type III.

To ensure that differences in parasite dissemination to the CNS were not driving the immune response differences, we quantified the parasite burden in the spleen, liver, lungs, and CNS at 5, 10, and 21 dpi using B1 qPCR. At these time points, we found that the parasite burden in these different organs did not differ between type II and type III-infected mice (Fig. 2C-F).

These data suggest that the CNS immune response differences we identified in type II and type III infection are not secondary to major differences in parasite dissemination to or persistence in the CNS, at least up to 21 dpi.

### Type III-infected mice have fewer alternatively activated macrophages and regulatory T cells in the CNS and spleen at 21 dpi

While our immunohistochemistry (IHC) data suggest that type III infection causes a higher number of macrophages and T cells to infiltrate into the CNS compared to type II infection, they do not address whether infection with type II or type III parasites affects the phenotype of these cells. To address this question, we isolated immune cells from the CNS of *Toxoplasma-*infected mice and then used flow cytometry to identify the frequency of different immune cell populations, focusing primarily on macrophages/microglia and T cells. Additionally, we performed the same studies on splenocytes from the infected mice to define if the CNS immune response was tissue-specific or merely reflective of differences in the global immune response.

As these studies represented the first studies to use flow cytometry to compare strain-specific CNS macrophage and T cell responses, we sought to profile major classes of cells by using previously identified markers (37–40). To this end, we focused on classically activated macrophages (M1s), alternatively activated macrophages (M2s), effector T cells (Teffs), and regulatory T cells (Tregs). The gating schemes we used are shown in **Fig. S1** (macrophages) **and Fig. S2** (T cells). In our analyses, we placed CD80/CD86 (M1s) and MMR/CXCR3 (M2s) in the same channels because transcriptional data have shown that M1s consistently co-express CD80 and CD86 (38) and a prior study that examined CNS macrophages in type II-infected mice showed that macrophages that express MMR also express CXCR3 (41). To validate this staining protocol, we verified that we obtained the same results regardless of whether CD80 and CD86 or CXCR3 and MMR are placed in individual channels or in the same channels (**Fig S3**). Finally, to further confirm the identity of the M1/M2 populations we isolated CD80^+^/CD86^+^ or MMR^+^/CXCR3^+^ splenocytes and used qPCR to determine the expression of IL-12, Arg-1, and IL-4 (21,22,39,41). As expected, the population we defined as M1s (CD80^+^/CD86^+^) expressed IL-12 and not Arg-1 and IL-4, while the M2s (MMR^+^/CXCR3^+^) expressed Arg-1 and IL-4 but not IL-12 (**Fig. S4A-D**). In addition, approximately 35% of our CD80^+^/CD86^+^ population was positive for IFN-γ while less than 1% of the MMR^+^/CXCR3^+^ population was positive for IFN-γ (**Fig. S4E,F**). Collectively, these data highly suggest that the macrophage population identified by CD45^+^, F4/80^+^, CD11b^hi^, CD11c^lo^, CD80^+^/CD86^+^ staining is consistent with prior descriptions of M1 macrophages and is pro-inflammatory. Similarly, the macrophage population identified by CD45^+^, F4/80^+^, CD11b^hi^, CD11c^lo^, MMR^+^/CXCR3^+^ staining is consistent with prior descriptions of M2 macrophages, which are less inflammatory. From here forward, we will simply denote these populations as M1s and M2s.

Based upon these validations, our flow analyses of the CNS immune cells showed that type III-infected mice had approximately half the number and frequency of M2s compared to type II-infected mice (Fig. 3A,B). For M1s, we observed no significant difference in the absolute number or frequency between the groups (Fig. 3C,D). In addition, we found that type III-infected mice had half the number and frequency of Tregs (CD3^+^, CD4^+^, FoxP3^+^) as compared to type II-infected mice (Fig. 3E,F) and no difference in the number or frequency of Teffs (CD3^+^, CD4^+^ or CD8^+^, CD44^+^) (**Table S2**).

**Fig 3.**
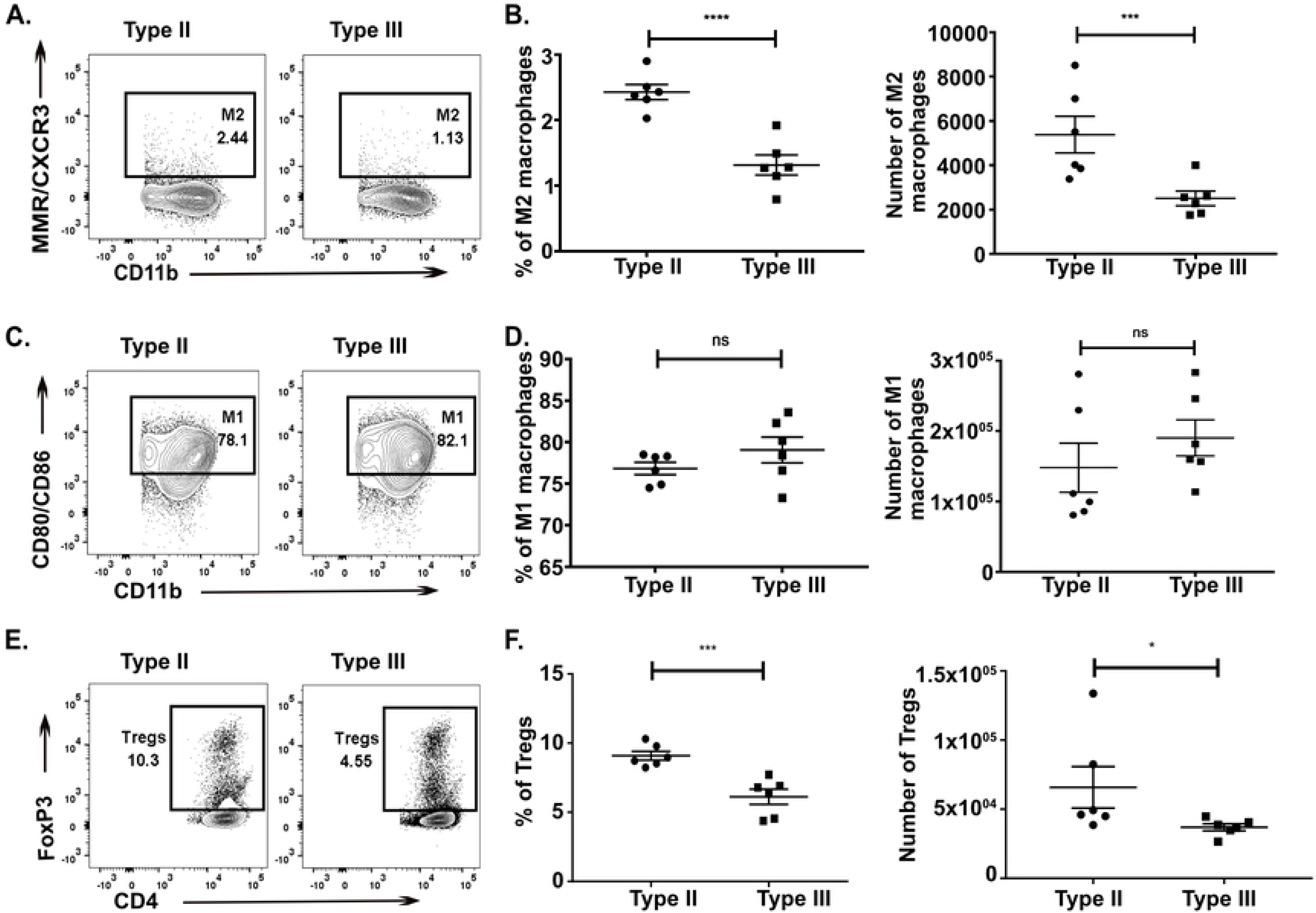
Type III-infected mice have fewer alternatively activated macrophages (M2) and T regulatory cells (Tregs) in the CNS as compared to type II-infected mice. At 21 dpi, immune cells were isolated from the CNS of either type II or type III-infected mice, split in half, and then stained for either T cell or macrophage markers. The stained cells were then analyzed by flow cytometry. **A,B.** CNS mononuclear cells evaluated for the presence of M2 macrophages (CD45^+^, F4/80^+^, CD11b^hi^, CD11c^lo^, MMR^+^/CXCR3^+^). **C,D.** CNS mononuclear cells evaluated for the presence of M1 macrophages (CD45^+^, F4/80^+^, CD11b^hi^, CD11c^lo^, CD80^+^/CD86^+^). **E,F.** CNS mononuclear cells evaluated for the presence of Tregs (CD3^+^ CD4^+^ FoxP3^+^). Bars, mean ± SEM. N= 6 mice/infected group. *p<0.05, **p<0.01, ***p<0.001, ****p<0.0001, ns= not significant, two-way ANOVA with Fisher’s protected LSD. Data representative of 2 individual experiments.

Consistent with our findings in the CNS we observed that splenocytes from type III-infected mice had approximately half the absolute number and frequency of M2s as compared to splenocytes from type II-infected mice (Fig. 4A,B). There was no significant difference in the absolute number and frequency of M1s in the spleen (Fig. 4C,D). Akin to our CNS data, splenocytes from type III-infected mice had half the number and frequency of Tregs compared to splenocytes from type II-infected mice (Fig. 4E,F).

**Fig 4.**
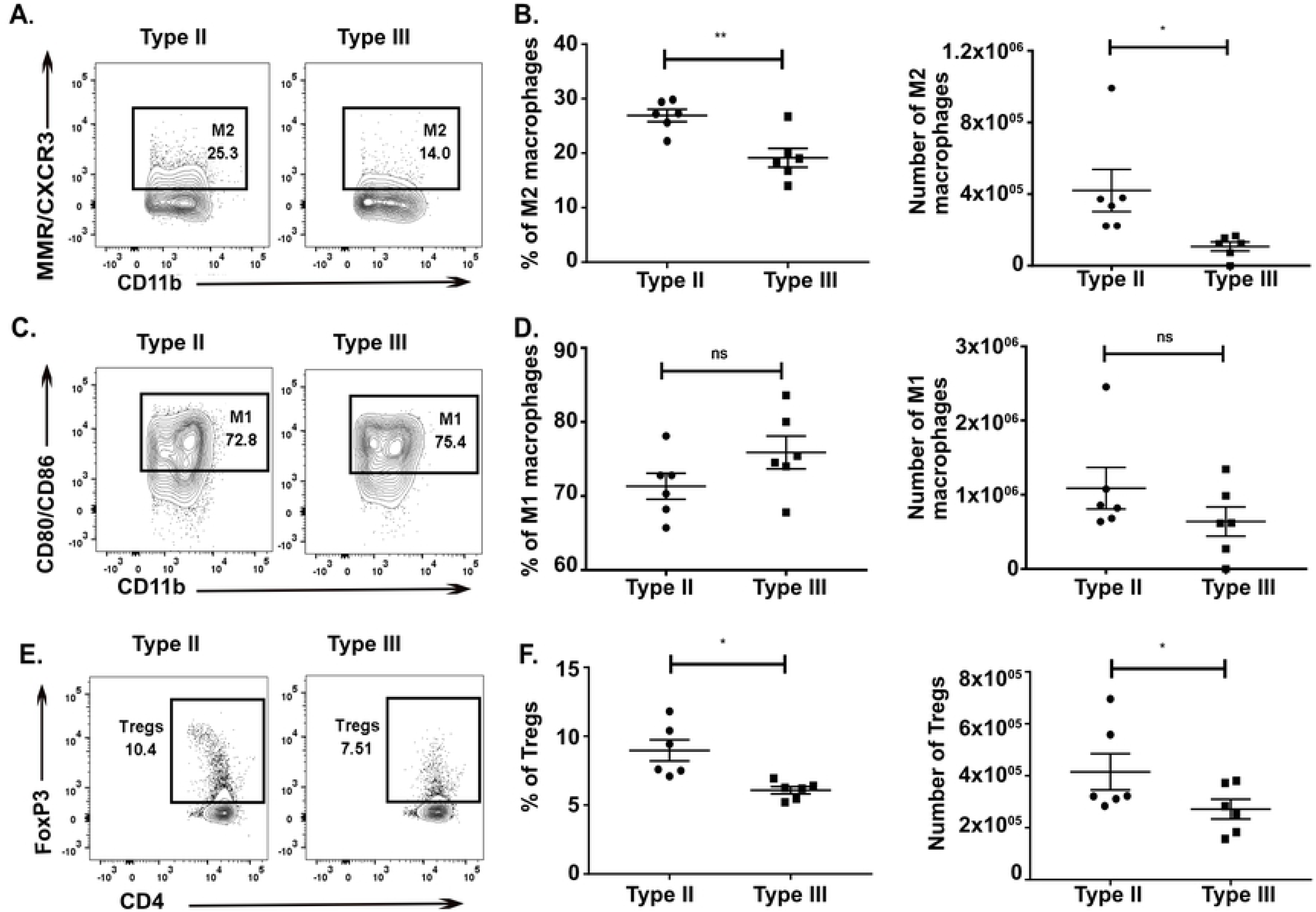
Type III-infected mice have fewer splenic alternatively activated macrophages (M2) and T regulatory cells (Tregs) compared to type II-infected mice. At 21 dpi, immune cells were isolated from the spleen of either type II- or type III-infected mice and stained and analyzed as in Fig.3. **A,B.** Splenic mononuclear cells evaluated for the presence of M2 macrophages. **C,D.** Splenic mononuclear cells evaluated for the presence of M1 macrophages. **E,F.** Splenic mononuclear cells evaluated for the presence of Tregs. Bars, mean ± SEM. N= 6 mice/infected group. *p<0.05, **p<0.01, ***p<0.001, ****p<0.0001, ns= not significant, two-way ANOVA with Fisher’s protected LSD. Data representative of 2 individual experiments.

The numbers of total CD3^+^ T cells, CD4 T cells (CD3^+^, CD4^+^), CD8 T cells (CD3^+^, CD8^+^), exhausted T cells (CD3^+^, CD8^+^, PD-1^+^), or macrophages (CD45^+^, F4/80^+^, CD11b^hi^, CD11c^lo^) in either the CNS or the spleen (**Table S2**) were not statistically different. In addition, to allow us to track infected cells and/or cells injected with parasite effector proteins (42), we infected Cre reporter mice that only express GFP after Cre-mediated recombination (43) with mCherry^+^ parasite strains that trigger Cre-mediated recombination (42,44). At 21 dpi, we identified no mCherry^+^ and/or GFP^+^ cells in the T cell or macrophage populations isolated from the CNS or spleen. The lack of GFP^+^ immune cells in the CNS at 21 dpi is consistent with our prior work using the same system (45).

These data strongly suggest that in addition to quantitative differences in the CNS immune response, type II and III-infected mice show differences in the phenotype of immune cells infiltrating into the CNS. Type III infection provokes a more pro-inflammatory sub-acute CNS immune response with a relative decrease in the macrophages (M2s) and T cells (Tregs) that suppress the pro-inflammatory response. In addition, as these differences are also seen in splenocytes, these data suggest that, at 21 dpi, the CNS immune response is reflective of the systemic immune response. Finally, the lack of mCherry^+^ and/or GFP^+^ immune cells suggests that our findings are not driven by a small population of immune cells that are actively infected or directly manipulated by parasites.

### The macrophage phenotype switches over time during type II and type III infection

Our finding that type III infection provokes a stronger pro-inflammatory response at 21 dpi was unexpected because of the prior work showing that macrophages infected with type III parasites are polarized to M2s while macrophages infected with type II parasites are polarized to M1s (21,46). As our work was done *in vivo* at 21 dpi and the prior work was done *in vitro* or very early *in vivo* (1-3 dpi), one explanation for these discrepancies is that the *in vivo* immune response evolves over time. To test this possibility and as we had found that splenocytes were accurate predictors of the CNS immune response, we phenotyped splenocytes from infected mice at 5, 10, and 21 dpi.

At 5 dpi, we observed that type III-infected mice had an approximately 3-fold higher frequency and 2-fold higher number of splenic M2s as compared to type II-infected mice (Fig. 5A). Conversely, at this time point, type II-infected mice showed an increased frequency and 1.5-fold higher number of splenic M1s as compared to type III-infected mice (Fig. 5B). By 10 dpi, the macrophage compartment from both type II and type III-infected mice had expanded and no difference in macrophage polarization state was seen (Fig. 5C-F). By 21 dpi, the splenic macrophage compartment was contracting and now type III-infected mice had fewer splenic M2s both by absolute number and frequency compared to type II-infected mice (Fig. 5C,D). At 5 and 10 dpi, for macrophages from type II or type III-infected mice, we found 1% or less of the macrophage population was infected or injected with parasite protein (i.e. ≤ 1% of the macrophage population was mCherry^+^ and/or GFP^+^). We found no strain-specific differences in the Tregs response at 5 or 10 dpi (Fig. 5G,H).

**Fig 5.**
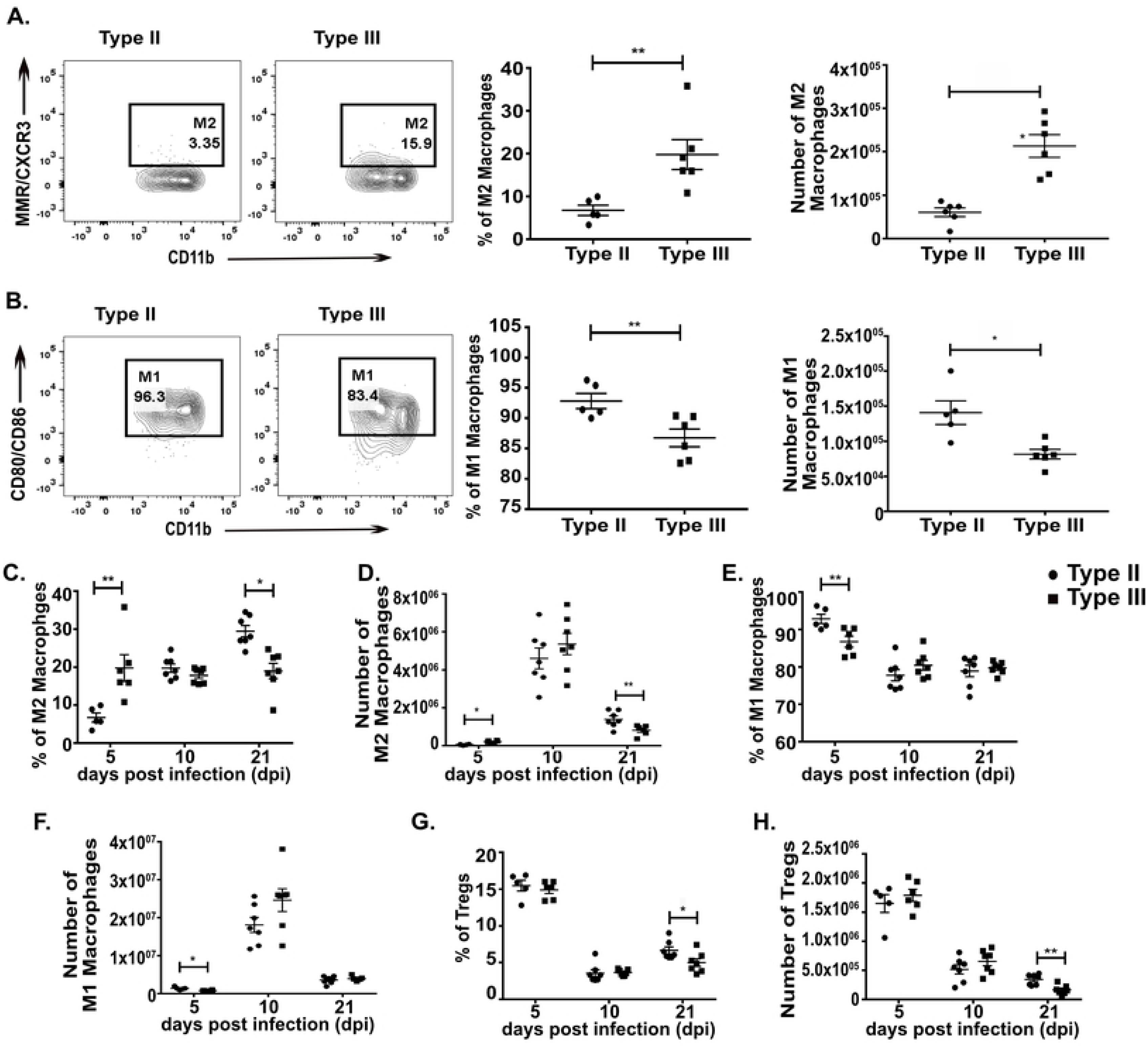
Macrophage phenotypes change over time during type II and type III infection. At 5, 10, and 21 dpi, immune cells were isolated from the spleen of either type II or type III-infected mice and stained and analyzed as in Fig.3. **A,B.** Quantification of the frequency and number of M2 **(A)** or M1 **(B)** macrophages at 5 dpi. **C,D.** Quantification of the frequency **(C)** and number **(D)** of splenic M2 macrophages over time. **E,F.** Quantification of the frequency **(E)** and number **(F)** of splenic M1 macrophages over time. **G,H.** Quantification of the frequency **(G)** and number **(H)** of splenic Tregs over time. Bars, mean ± SEM. N= 5-7 mice/infected group. *p<0.05, **p<0.01, two-way ANOVA with Fisher’s protected LSD. Data representative of 2 individual experiments. • = type II, ■ = type III.

These data show that early in infection type III-infected mice show a stronger M2 macrophage response than type II-infected mice. However, as the infection progresses, the immune response evolves such that by 21 dpi type III-infected mice now have a significant decrease in these anti-inflammatory macrophages compared to type II-infected mice, even though parasite dissemination to the CNS is equivalent (Fig. 2). Unlike the macrophage response, we did not observe strain-specific differences in Tregs until 21 dpi. Our data also show that these strain-specific differences are primarily found in uninfected macrophages.

### *ROP16_III_* affects the type III CNS immune response and parasite persistence

Given the evolution of these strain-specific differences from 5 to 21 dpi, we hypothesized that the early macrophage immune response might heavily influence the ensuing sub-acute immune response. We focused on the early macrophage response for several reasons. Tissue resident macrophages are some of the first cells to interface and respond to infecting microbes (47). Consistent with this concept, at 5 dpi, we found strain-specific differences in the macrophage compartment but not the T cell compartment (Fig. 5A-H). Furthermore, a growing body of literature suggests that this early response influences the ensuing T cell response, possibly through the secretion of cytokines (48). Finally, as noted above, prior *in vitro* and *in vivo* work has already established that macrophages infected with type II parasites polarize to M1s while macrophage infected with type III parasites polarize to M2s. Importantly, these studies also determined that specific alleles of *Toxoplasma* effector proteins (GRA15_II_ for M1s and ROP16_III_ for M2s) drive these macrophage phenotypes (19,21,22), giving us a mechanism for altering these responses. Thus, to determine if early macrophage responses influence the subsequent CNS immune response, we used CRISPR/Cas9 (49–53) to engineer a type III strain that lacked *rop16* (**Fig. S5A**). We validated the deletion of *rop16* (IIIΔ*rop16*) using locus-specific PCR (**Fig. S5B**) and a functional assay to show that these parasites no longer induced host cell phosphorylation of STAT6 (**Fig. S5C**), the transcription factor linked to the *rop16*_III_-associated M2 phenotype (19). Given that this strain should lack the early type III-associated M2 response, we predicted that it would provoke a more type II-like sub-acute CNS immune response. To test this prediction, we infected mice with type II, type III, or IIIΔ*rop16* parasites and, at 21 dpi, analyzed the CNS immune response by IHC and flow cytometry. Consistent with our hypothesis, by quantitative IHC, we found that the CNS immune response in IIIΔ*rop16*-infected mice looked akin to type II-infected mice with fewer infiltrating macrophages/microglia and T cells compared to type III-infected mice (Fig. 6A-E). By flow cytometry, IIIΔ*rop16*-infected mice again looked similar to type II-infected mice in terms of M2s frequency and absolute number (Fig. 7A,B). The frequency and the absolute number of M1s were not statistically different between type II, type III, or IIIΔ*rop16*-infected mice (Fig. 7C,D). Unexpectedly, by both *Toxoplasma*-specific qPCR and cyst count, the IIIΔ*rop16*-infected mice showed a substantial decrease in the CNS parasite burden compared to type II or type III-infected mice (Fig. 8A,B).

**Fig 6.**
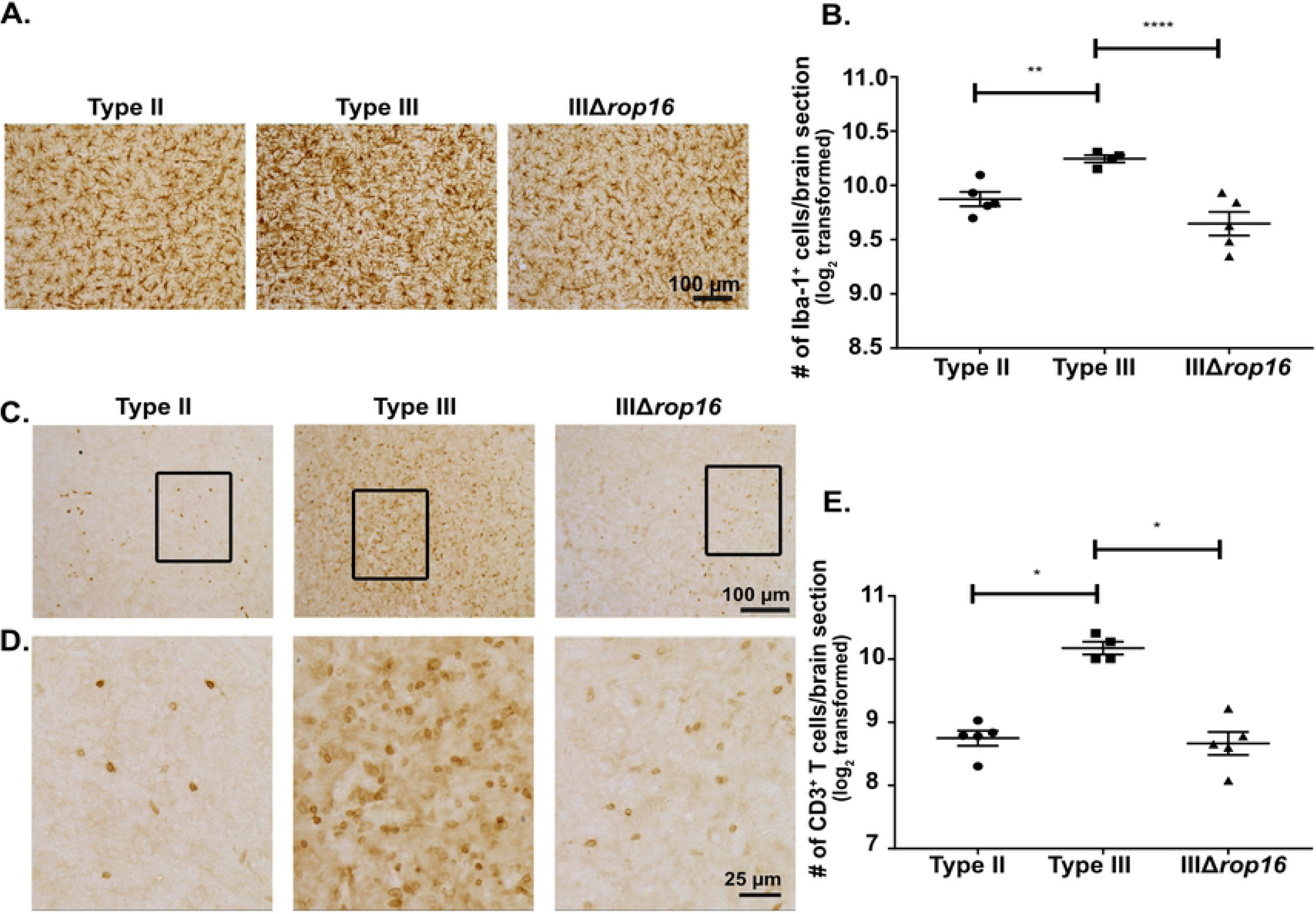
IIIΔ*rop16*-infected mice show a type II-like CNS response with fewer CNS macrophages/microglia and T cells. Mice were inoculated with type II, type III, or IIIΔ*rop16* parasites, and brains harvested at 21 dpi. Macrophages/microglia and T cells infiltration into the CNS was determined by quantitative IHC (as in Fig.1). **A.** Representative image of Iba-1^+^ cells (macrophage/microglia). **B.** Quantification of the number of Iba-1^+^ cells. **C.** Representative image of CD3^+^ cells (T cell). **D.** Enlargement of the boxed area in **(C)**. **E.** Quantification of the number of CD3^+^ cells. Bars, mean ± SEM. N= 12 fields of view/section, 3 sections/mouse, 5 mice/infected group. For each mouse, the number of cells/section was averaged to create a single point. *p<0.05, two-way ANOVA with Fisher’s protected LSD. Data representative of 3 individual experiments with two different, individually engineered IIIΔ*rop16* clones.

**Fig 7.**
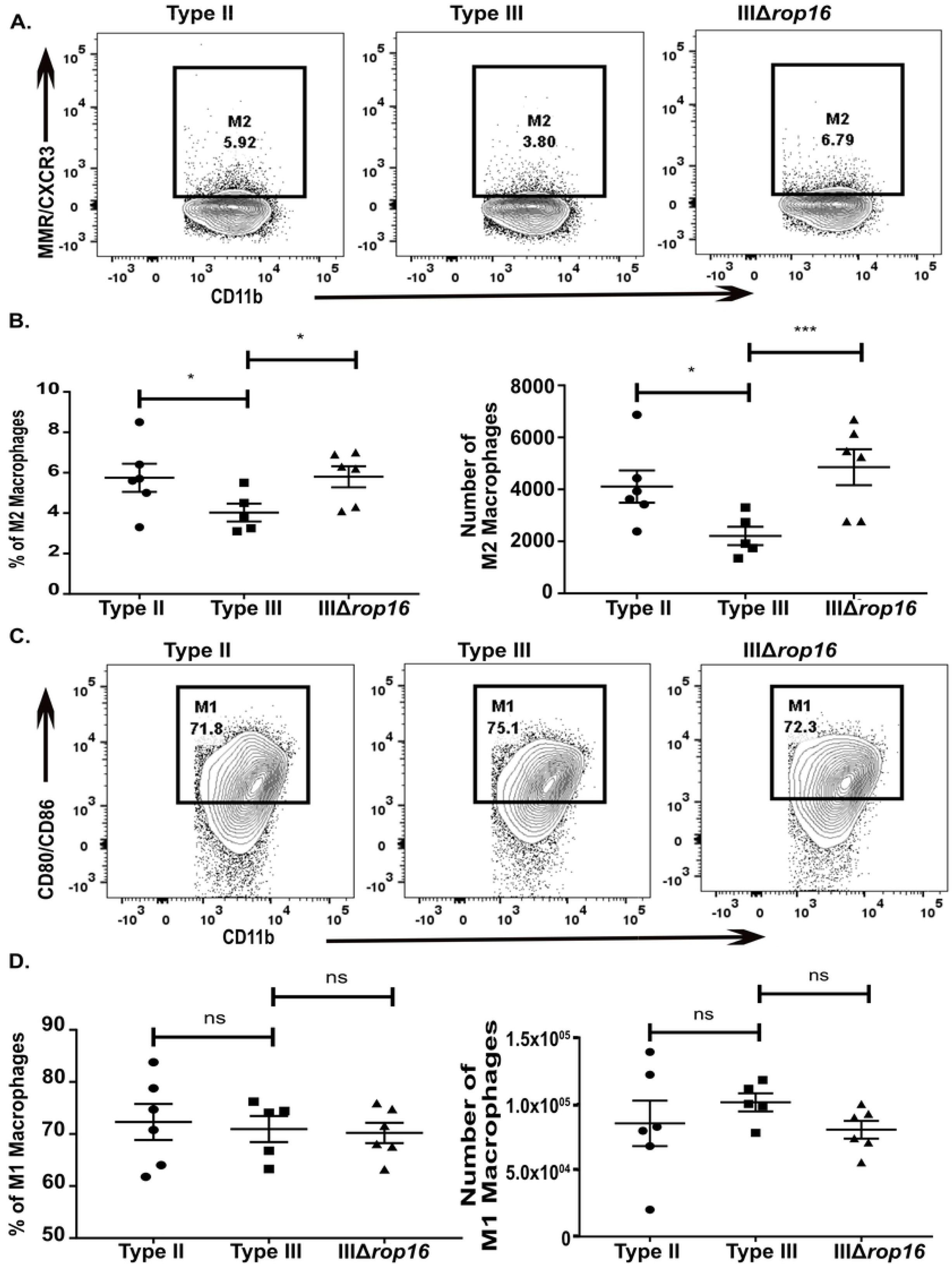
IIIΔ*rop16* infection provokes more alternatively activated macrophages, akin to a type II CNS immune response. At 21 dpi, immune cells were isolated from brains infected mice and then stained and analyzed as in Fig.3. **A,B.** CNS mononuclear cells evaluated for the presence of M2 macrophages. **C,D.** CNS mononuclear cells evaluated for the presence of M1 macrophages. Bars, mean ± SEM. N= 4-5 mice/infected group. *p<0.05, **p<0.01, ***p<0.001, two-way ANOVA with Fisher’s protected LSD. Data representative of 3 individual experiments with two different, individually engineered IIIΔ*rop16* clones.

**Fig 8.**
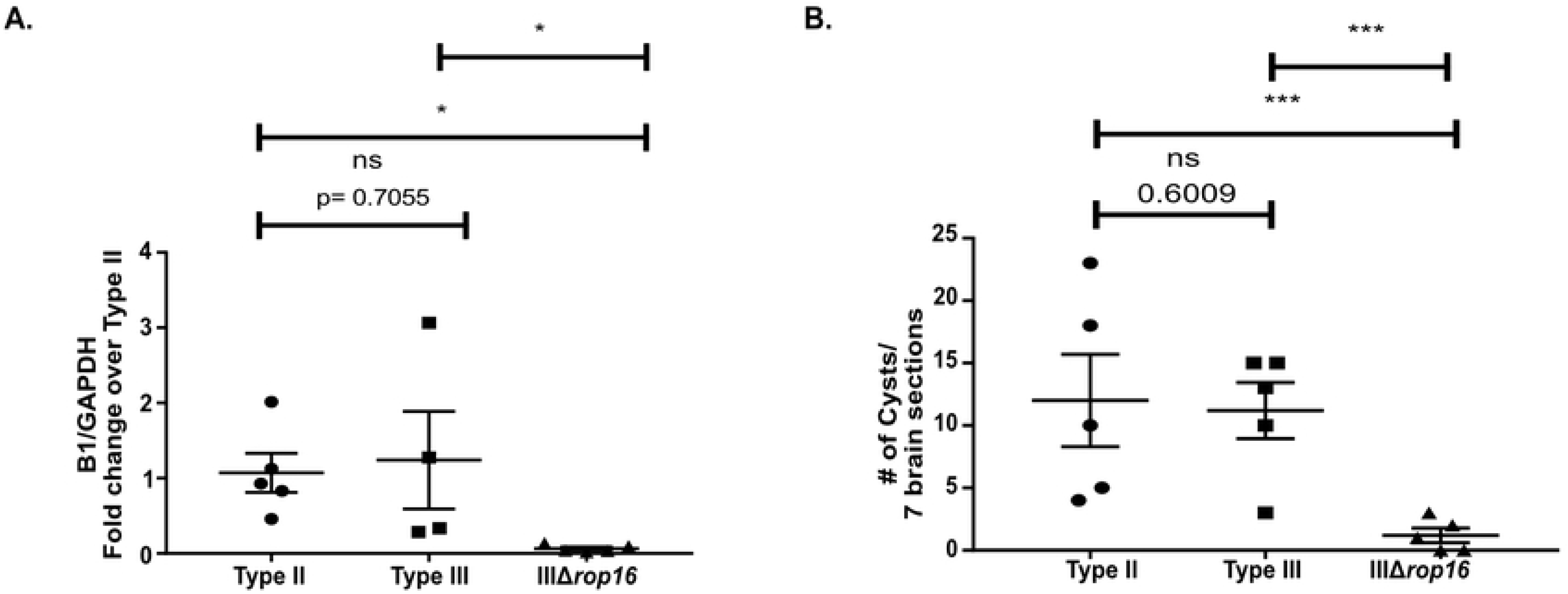
IIIΔ*rop16*-infected mice had a significantly lower CNS parasite burden than type II and type III-infected mice. Mice were inoculated with type II, type III, or IIIΔ*rop16* parasites. Brains were harvested at 21 dpi and analyzed as in Fig.2. **A.** Quantification of CNS *Toxoplasma* burden by qPCR for the *Toxoplasma*-specific B1 gene. **B.** Quantification of *Toxoplasma* cyst burden in brain sections stained with DBA. Bars, mean ± SEM. N= 5 mice/infected group. *p<0.05, ns=not significant, two-way ANOVA with Fisher’s protected LSD. Data representative of 3 individual experiments with two different, individually engineered IIIΔ*rop16* clones.

To verify that the lack of *rop16_III_* drove the immune response and parasite burden changes we identified, we generated a IIIΔ*rop16::ROP16_III_* strain that ectopically expresses *rop16_III_*. We validated the expression of *rop16_III_* using gene-specific PCR and a functional assay to confirm parasite-induced host cell phosphorylation of STAT6 (**Fig. S5A-C**). We then infected mice with type III, IIIΔ*rop16*, or IIIΔ*rop16::ROP16_III_* parasites and analyzed the CNS immune response by quantitative IHC and flow cytometry. The IIIΔ*rop16::ROP16_III_* strain produced a CNS immune response akin to the parental type III strain, and distinct from the IIIΔ*rop16*, in terms of macrophage and T cell numbers (Fig. 9A,B), parasite burden (Fig. 9C,D), and M2 number and frequency (Fig. 9E). The M1 number and frequency was not different between the three strains (Fig. 9F).

**Fig 9.**
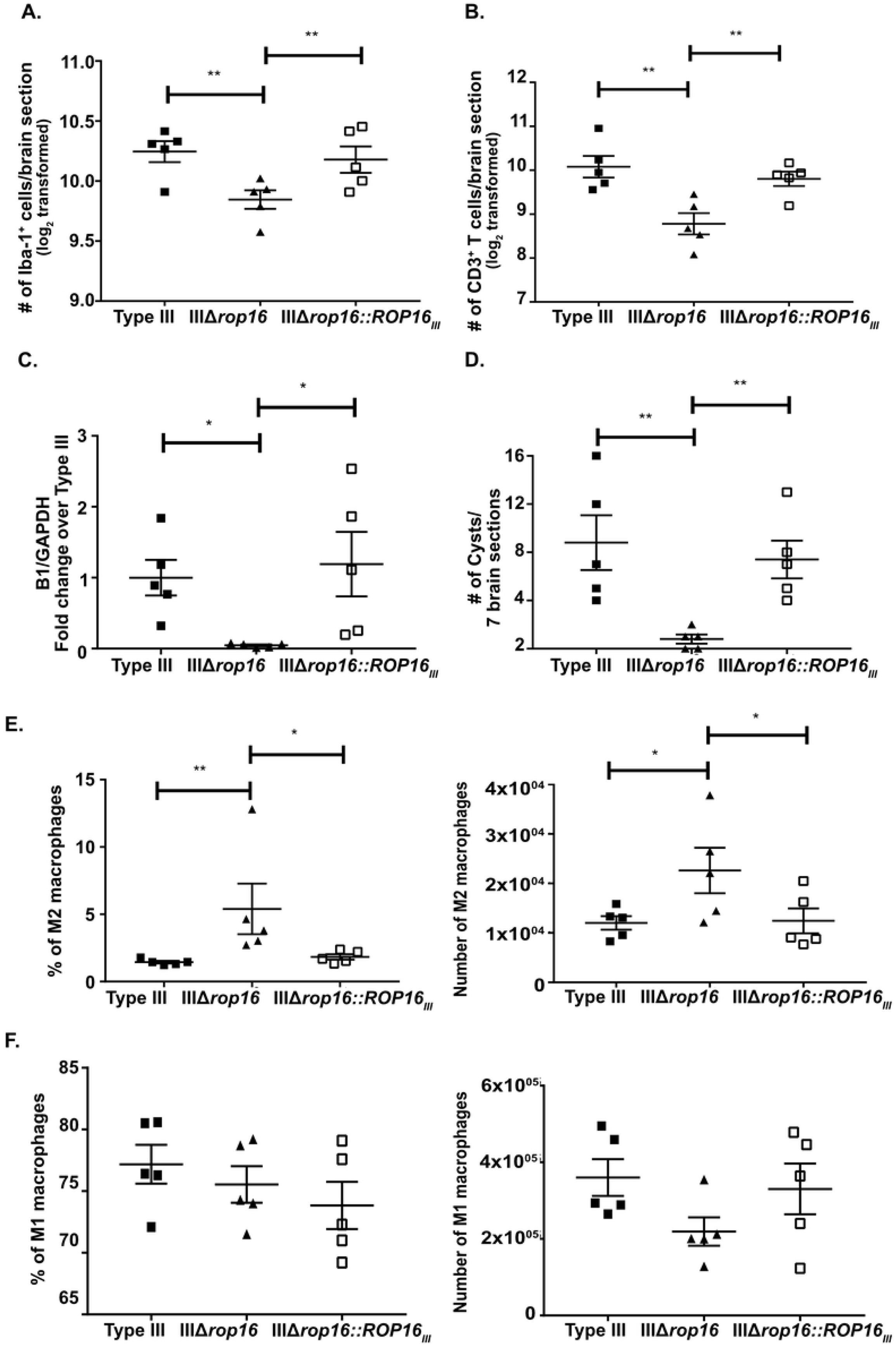
The infection with IIIΔ*rop16::ROP16_III_* parasites restores type III-like CNS response with higher CNS macrophages/microglia and T cells. Mice were inoculated with type III, IIIΔ*rop16*, or IIIΔ*rop16:ROP16* parasites. Brains were harvested at 21 dpi and analyzed as in Fig 1 (**A,B**) or Fig 2 (**C,D**). **A.** Quantification of the number of Iba-1^+^ cells (macrophages/microglia). **B.** Quantification of the number of CD3^+^ T cells. Bars, mean ± SEM. N= 12 fields of view/section, 3 sections/mouse, 4-5 mice/infected group. For each mouse, the number of cells/section was averaged to create a single point. **C.** Quantification of CNS *Toxoplasma* burden by qPCR for the *Toxoplasma*-specific B1 gene. **D.** Quantification of *Toxoplasma* cyst burden in brain sections stained with DBA. At 21 dpi, immune cells were isolated from brains infected mice and then stained and analyzed as in Fig.3. **E.** CNS mononuclear cells evaluated for the presence of M2 macrophages. **F.** CNS mononuclear cells evaluated for the presence of M1 macrophages. Bars, mean ± SEM. N= 12 fields of view/section, 3 sections/mouse, 4-5 mice/infected group. *p<0.05, **p<0.01, two-way ANOVA with Fisher’s protected LSD. Data representative of 2 individual experiments using a single IIIΔ*rop16* clone and IIIΔ*rop16:ROP16* clone.

In summary, these data highly suggest that, in the context of a type III infection, *rop16_III_* influences the CNS immune response and enables parasite persistence.

### A lack of *rop16_III_* induces a mixed inflammatory response during acute infection

To confirm that the changes in the CNS were downstream of a change in the acute inflammatory response, at 5 dpi we phenotyped splenocytes from mice infected with type II, type III, or IIIΔ*rop16* parasites. Unexpectedly, mice infected with the IIIΔ*rop16* strain showed a mixed immune phenotype with an increase in both M2s and M1s (Fig. 10A,B). In addition, and unlike either type II or type III-infected mice, IIIΔ*rop16*-infected mice showed an increase in the number and frequency of Tregs (Fig. 10C) as well as a 3-fold increase in the number of IFN-γ^+^ CD4 and CD8 T cells (Fig. 10D,E). Using the mean fluorescence intensity of IFN-γ staining to assess IFN-γ production per cell, we found that T cells from IIIΔ*rop16* and type II-infected mice produced similar amounts of IFN-γ, which was 1.5-fold higher than T cells from type III-infected mice (Fig. 10D,E).

**Fig 10.**
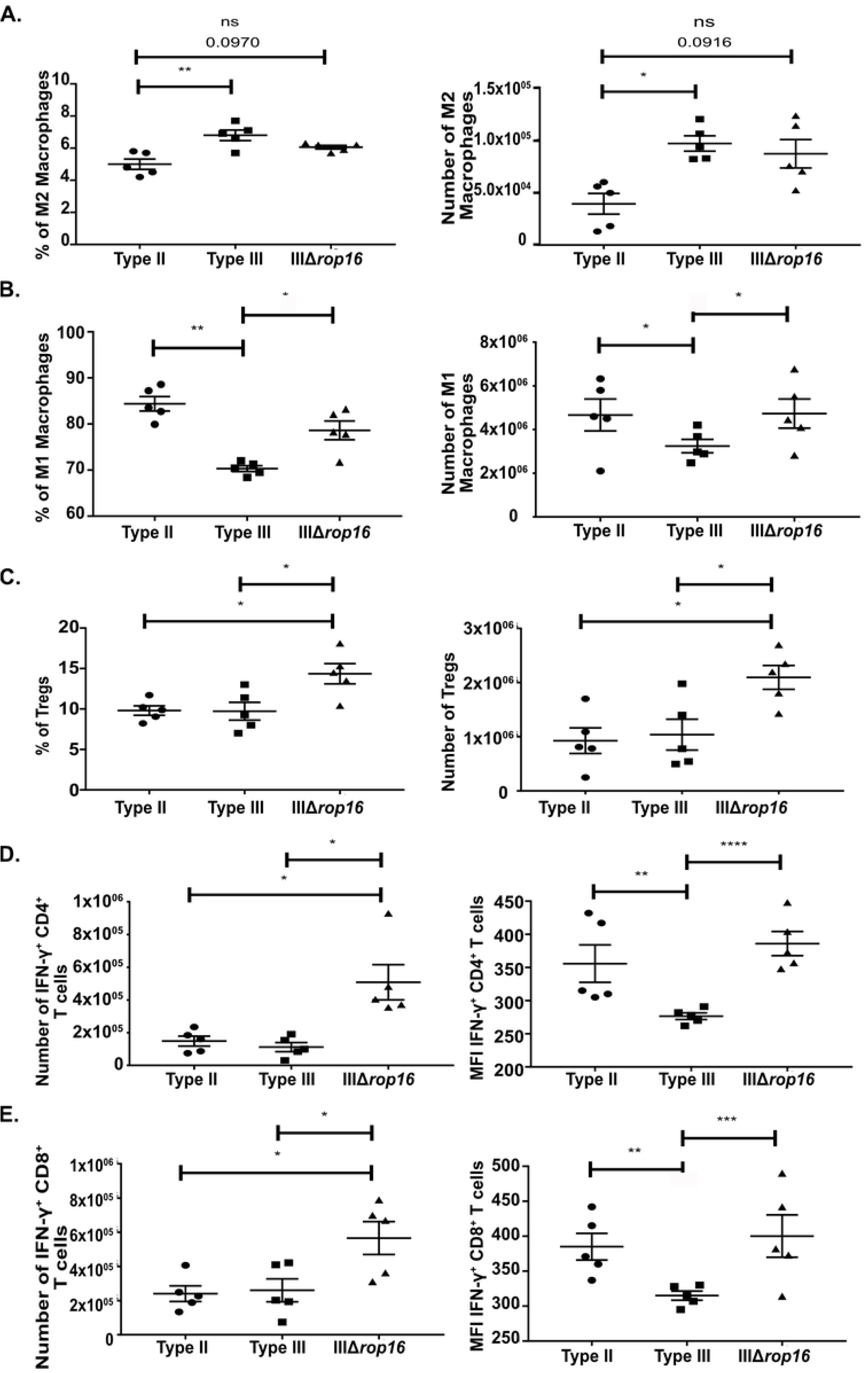
IIIΔ*rop16* infected mice showed a mixed immune response in the periphery. At 5 dpi, immune cells were isolated from the spleen of infected mice and then stained and analyzed as in Fig. 3. **A.** Quantification of the frequency and number of splenic M2 macrophages. **B.** Quantification of the frequency and number of splenic M1 macrophages. **C.** Quantification of the frequency and number of Tregs. **D.** Quantification of the number and mean fluorescence intensity of splenic IFN-γ producing CD4^+^ T cells. **F.** Quantification of the number and mean fluorescence intensity of splenic IFN-γ producing CD8^+^ T cells. Bars, mean ± SEM N= 5 mice/infected group. *p<0.05, **p<0.01, ***p<0.001, ****p<0.0001, two-way ANOVA with Fisher’s protected LSD. Data representative of 3 individual experiments with two different, individually engineered IIIΔ*rop16* clones.

To verify that these acute peripheral immune response changes were driven by *rop16_III_*, we infected mice with the type III, IIIΔ*rop16*, or IIIΔ*rop16::ROP16_III_* parasites and phenotyped the splenocytes at 5 dpi. As expected the IIIΔ*rop16::ROP16_III_* strain produced a splenocyte immune response akin to the parental type III strain in terms of M2s, M1s, and Treg frequency and absolute number (Fig. 11A-C). In the IFN-γ^+^ CD4 and CD8 compartment, the IIIΔ*rop16::ROP16_III_* strain also produced a parental type III response in terms of absolute numbers of CD4 and CD8 IFN-γ^+^ cells and the level of IFN-γ produced per cell (Fig. 11D,E).

**Fig 11.**
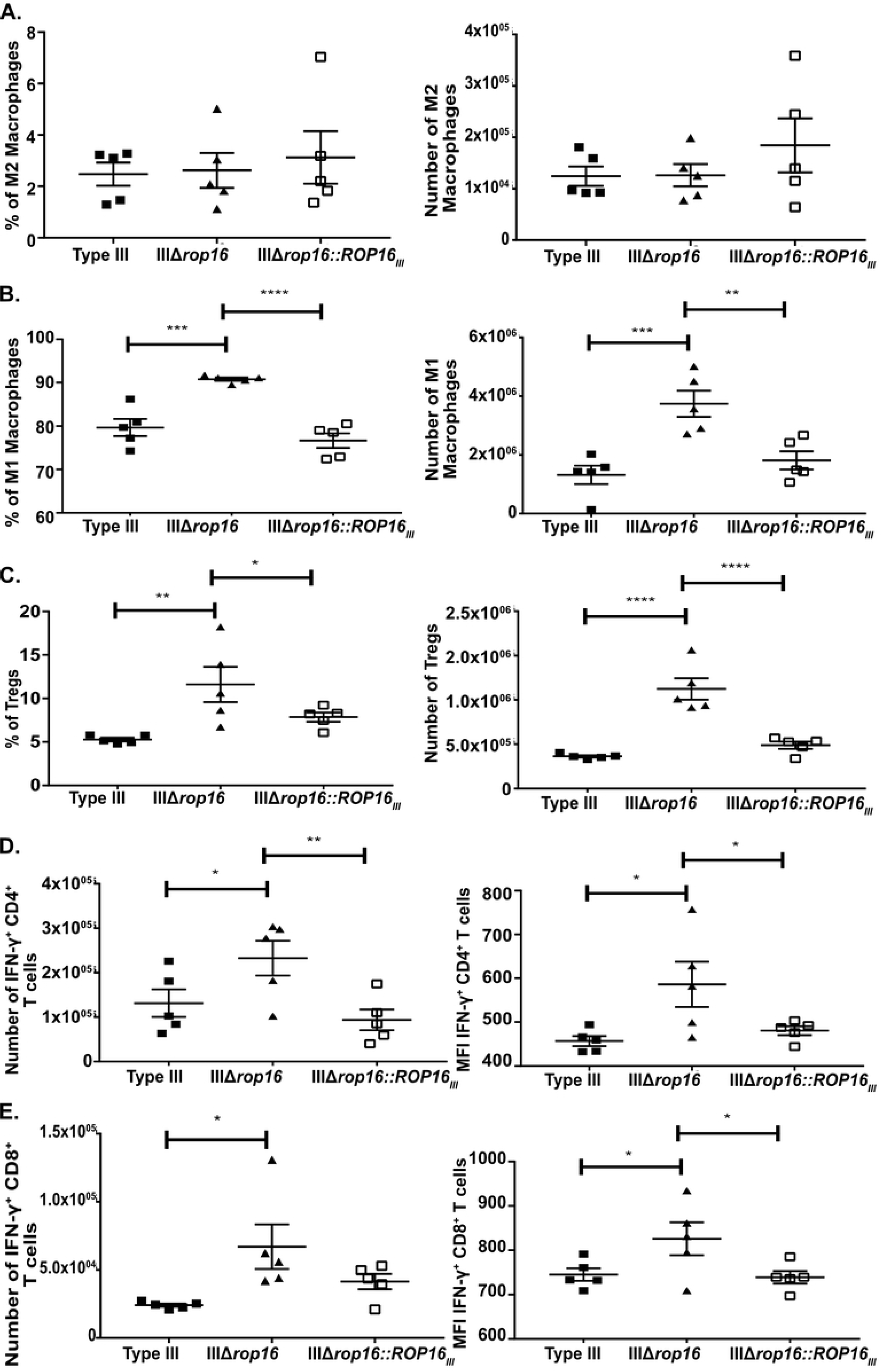
IIIΔ*rop16::ROP16_III_* infected mice showed a type III-like immune response in the periphery. At 5 dpi, immune cells were isolated from the spleen of infected mice and then stained and analyzed as in Fig 3. **A.** Quantification of the frequency and number of splenic M2 macrophages. **B.** Quantification of the frequency and number of splenic M1 macrophages. **C.** Quantification of the frequency and number of splenic Tregs. **D.** Quantification of the number and mean fluorescence intensity of splenic IFN-γ producing CD4^+^ T cells. **E.** Quantification of the number and mean fluorescence intensity of splenic IFN-γ producing CD8^+^ T cells. Bars, mean ± SEM N= 5 mice/infected group. *p<0.05, **p<0.01, ***p<0.001, ****p<0.0001, two-way ANOVA with Fisher’s protected LSD. Data representative of 2 individual experiments using a single IIIΔ*rop16* clone and IIIΔ*rop16:ROP16* clone.

In summary, these data show that type III parasites lacking *rop16* produce a mixed inflammatory phenotype at 5 dpi, suggesting that *rop16_III_* is an important determinant of the acute immune response to type III parasites.

### Type III parasites depend upon *rop16* to avoid early clearance by Immunity-Related GTPases (IRGs)

Given this unanticipated acute inflammatory response including the elevated level of T cell IFN-γ production, and decreased CNS parasite burden at 21 dpi, we hypothesized that the IIIΔ*rop16* parasites were undergoing an increased level of early clearance. This heightened early inflammatory response and the rapid clearance of parasites would then provoke a strong counterbalancing anti-inflammatory response, causing an increase in both M2s and Tregs early in infection. This hypothesis is particularly appealing because type III strains have a very low expression of the virulent allele of *rop18*, a parasite virulence gene that disables the interferon-γ-inducible Immunity-Related GTPase system (IRGs) which is a major mechanism by which murine host cells kill intracellular parasites in an IFN-γ dependent manner (25,26,54). To test this possibility, we quantified the frequency of infected peritoneal exudate cells (PECs) at 1, 3, and 5 dpi from mice infected with type II, type III, or IIIΔ*rop16* parasites. At 1 and 3 dpi, all 3 strains showed the same frequency of infected PECs. But, by 5 dpi, IIIΔ*rop16*-infected mice showed a 4-6-fold lower rate of infected PECs compared to type II or parental type III-infected mice (Fig. 12A-C). To confirm that this increase in parasite clearance was secondary to a lack of *rop16_III_*, we infected mice with type III, IIIΔ*rop16*, or IIIΔ*rop16::ROP16_III_* parasites and assayed the frequency of infected PECs at 3 and 5 dpi. As expected, the IIIΔ*rop16::ROP16_III_* strain maintained parental levels of infected PECs at both time points (Fig. 12D).

**Fig 12.**
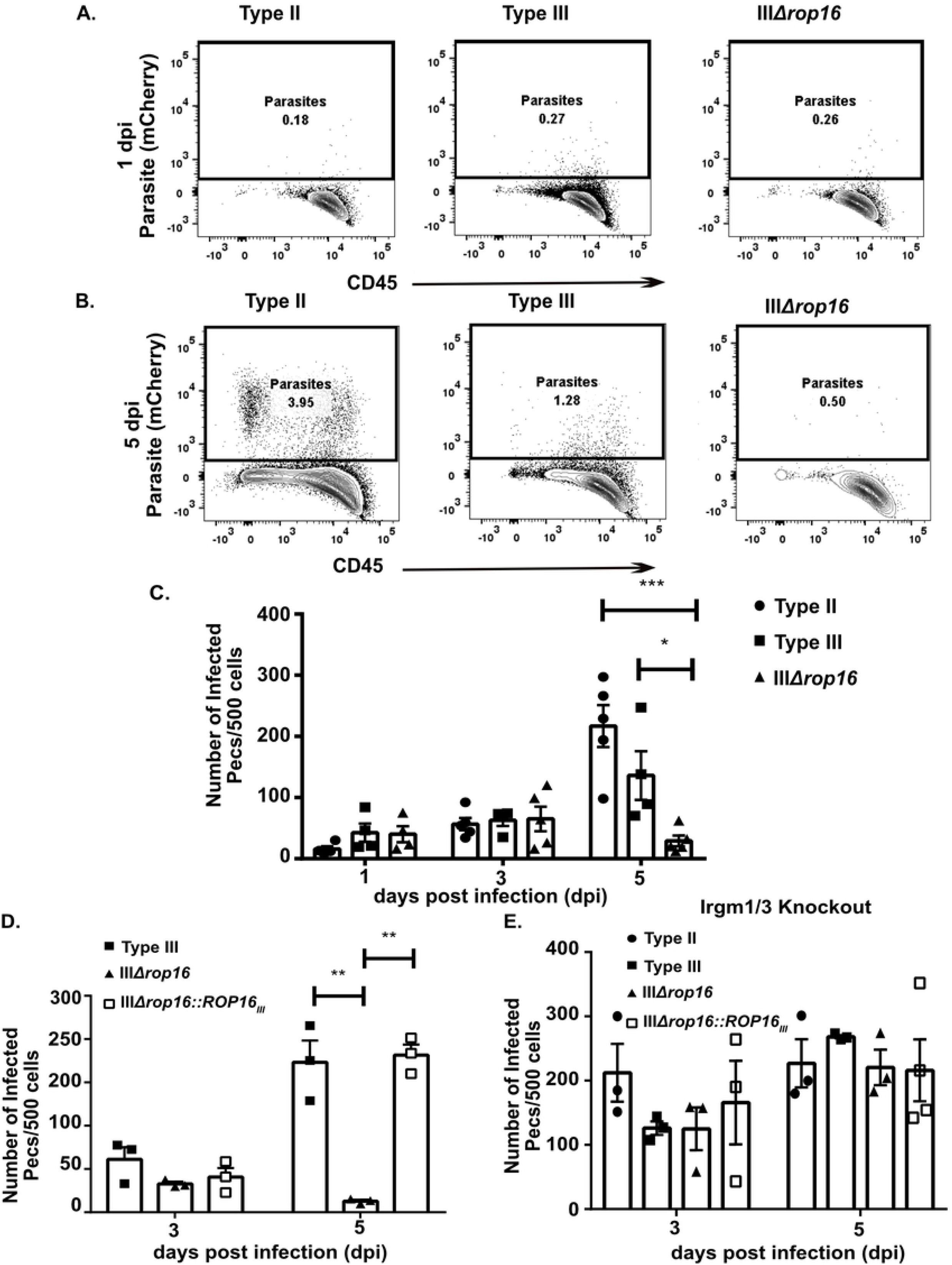
IIIΔ*rop16* parasites are cleared early *in vivo*. At 1, 3, and 5 dpi, peritoneal exudate cells (PECs) were isolated from infected mice, stained for CD45, and then screened for infected cells (CD45^+^, mCherry^+^). **A,B.** Representative plots of infected PECs at **(A)** 1 dpi and **(B)** 5 dpi. **C.** Quantification of the number of infected cells over time. Bars, mean ± SEM. N=5 mice/infected group. *p<0.05, ***p<0.001, two-way ANOVA with Fisher’s protected LSD. Data representative of 2 individual experiments with two different, individually engineered IIIΔ*rop16* clones. **D.** PECs isolated and quantified as in **(C)**. Quantification of the number of infected cells over time. **E.** PECs isolated and quantified as in **(C)** but using *Irgm1/3^-/-^* mice. Quantification of the number of infected cells over time. (**D,E)** Bars, mean ± SEM. N=3-4 mice/infected group. A single IIIΔ*rop16* clone and IIIΔ*rop16:ROP16* clone were used. **p<0.01, one-way ANOVA with Fisher’s protected LSD. • = type II, ■ = type III, ▲ = IIIΔ*rop16*, □ = IIIΔ*rop16::ROP16_III_*.

Finally, to directly test if the IRGs were the mechanism by which the IIIΔ*rop16* parasites were being cleared, we assayed the frequency of infected PECs at 3 and 5 dpi in mice that lack both *Irgm1* and *Irgm3*, key mediators of IRG pathway (27,55–58). We reasoned that if the IRGs mediated the *rop16_III_*-dependent increase in clearance, then in Irgm1/3 KO mice, the IIIΔ*rop16* strain should now maintain parental levels of PECs infection at 5dpi, which is what we found (Fig. 12E).

Collectively, these data suggest that, in the context of a type III infection, *rop16_III_* is essential to dampen the acute IFN-γ response thereby avoiding rapid parasite clearance by the IRGs.

## Discussion

The results presented here show that genetically distinct *Toxoplasma gondii* strains provoke strain-specific CNS immune responses and that these sub-acute immune responses are likely influenced by the initial systemic immune response. We have shown that, compared to infection with a type II strain, infection with a type III strain induces a more pro-inflammatory, sub-acute CNS immune response in both quality and quantity at the level of infiltrating T cells and macrophages/microglia, and that these strain-specific immune responses are not simply driven by differences in parasite burden. In addition, we have shown that, for the parameters monitored at 21 dpi, the CNS immune response mirrors the systemic immune response as gauged by splenocytes. By temporally profiling the systemic macrophage response, we have shown that this response evolves over time, leading us to hypothesize that the early macrophage response affects the subsequent sub-acute response. This hypothesis is partially supported by our finding that a IIIΔ*rop16* strain, which induces an early immune response distinct from the parental type III strain, produces a type II-like CNS immune response in quantity and quality, despite having a much lower CNS parasite burden than either the type II or parental type III strain.

Based upon these data and prior work, we propose the following model: early in infection, type II-infected macrophages are polarized to M1s which secrete high levels of IL-12 (21,22). This secreted IL-12 then influences the uninfected macrophages to polarize towards M1s, resulting in a systemic, highly pro-inflammatory early M1 responses (5 dpi), with high levels of IL-12 and IFN-γ production (**Fig S4**). This early pro-inflammatory response then provokes a counter-balancing anti-inflammatory response that leads to a rise in M2s, which continues as parasites disseminate and enter the brain. This systemic response is then recapitulated in the CNS as parasites establish a chronic CNS infection (Fig. 13A). Conversely, for type III parasites, during acute infection, type III-infected macrophages are polarized to M2s via *rop16_III_* leading to increased IL-4 production (**Fig S4**), which promotes a mixed systemic inflammatory response with more early M2s, resulting in decreased levels of IFN-γ and IL-12 and increased levels of IL-4 (**Fig S4**) compared to type II-infected mice. This less pro-inflammatory early response enables type III parasites to evade early clearance and avoids provoking the compensatory anti-inflammatory response, so as type III parasites disseminate to the CNS, the immune response that occurs is now strongly pro-inflammatory (Fig. 13B).

**Fig 13.**
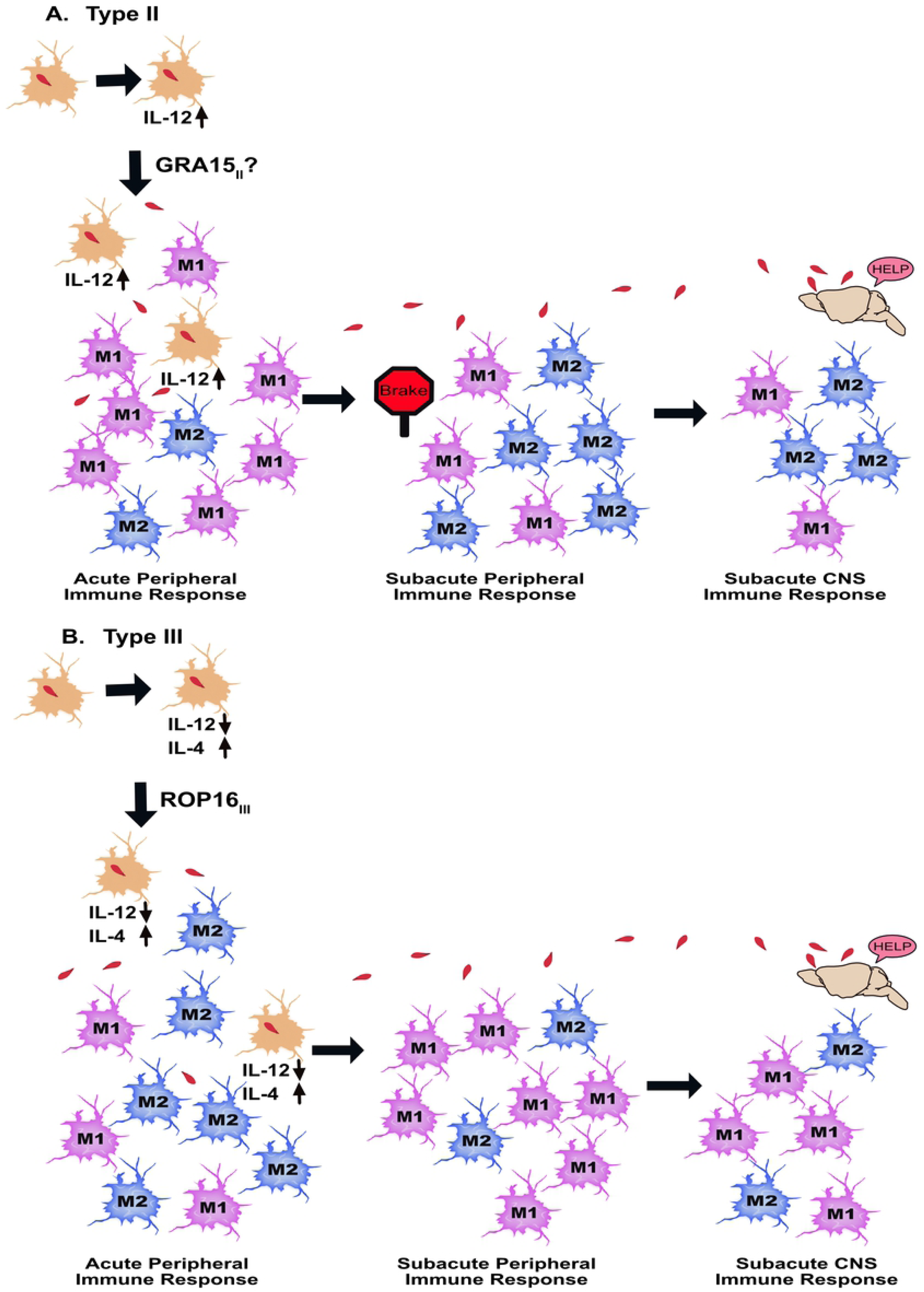
Model for strain-specific acute and sub-acute macrophage immune responses. **A.** During acute infection, type II parasites infect tissue resident macrophages and, via GRA15_II_, modulate these macrophages to polarize to M1 macrophages and secrete IL-12. This acute elevation of IL-12 leads to more uninfected macrophages polarizing to M1s, which will lead to more IFN-γ production by both macrophages and T effector cells (Teffs). This early pro-inflammatory immune response then initiates a compensatory anti-inflammatory immune response especially at the level of M2 macrophages. During this time, parasites proliferate and disseminate to the brain, at which point the immune cells that are present in the periphery infiltrate into the brain. **B.** During acute infection, the type III parasites infect tissue resident macrophages, and, via ROP16_I/III_, modulate these macrophages to polarize to M2 macrophages which produce less IL-12 and more IL-4. In turn, the increased IL-4 and decreased IL-12 leads to an increased level of uninfected M2 polarized macrophages and Teffs that produce less IFN-γ. This less pro-inflammatory response avoids the early compensatory anti-inflammatory immune response. As type III parasites proliferate and disseminate, a highly inflammatory response ensues, which then infiltrates into the brain.

While these models fit our data, many unanswered questions remain. What are the molecular and cellular mechanisms by which these early immune responses influence the later responses? How does the type II *gra15* allele, which drives an M1 phenotype in infected macrophages, influence the systemic immune response to type II strains? Do these differences persist even in highly established chronic infections (e.g. months to years post infection)? Though these strain-specific immune responses do not appear to be driven by gross differences in systemic dissemination or CNS parasite burden, could strain-specific rates of switching from tachyzoites to bradyzoites play a role in driving immune response differences? We believe we have established a system in which these highly complex host-parasite interactions can systematically be dissected using engineered parasites and mice.

In addition to establishing a tractable model for understanding the evolution of immune responses, several other important points arise from our data. We potentially identified a reason for the retention of the type I/III *rop16_I/III_* (*rop16_I/III_*) allele. In the original type II x III cross, *rop16* was not identified as a virulence gene but rather its strain specificity was detected through strain-specific differences in host cell signaling in human fibroblasts (59). In fact, in a highly lethal type I strain, at 72 hours post infection, the loss of *rop16* increased the PECs infection rate, systemic dissemination, and parasite burden in distal organs while also increasing IL-12 production by PECs. These data suggest that the increase in IL-12, which should result in higher IFN-γ production, does not adversely affect the IΔ*rop16* parasites (19). Conversely, we used a type III strain, which is genetically similar to the type I strain but avirulent in mice because of its’ low expression of *rop18*, a key protein for blocking murine IFN-γ-dependent cell intrinsic defenses (25,26,54). In the context of low *rop18* expression and therefore high susceptibility to the IRGs, the ability of *rop16_I/III_* to decrease the early IFN-γ response (Fig. 10E) by decreasing IL-12 and increasing the M2 response, appears crucial for type III strains to avoid rapid clearance during the very earliest part of infection. This proposed mechanism is supported by the data in the Irgm1/3 knockout mice (Fig. 12) as well as prior work showing that IRG-mediated clearance of intracellular parasites is a major murine IFN-γ-dependent mechanism for controlling *Toxoplasma* (25,26,54). Collectively, these data suggest that, *in vivo*, *rop16_I/III_* is dispensable for type I strains but essential for type III strains, a discrepancy potentially explained by differences in *rop18* expression.

We have also shown that the strain-specific polarizations previously identified in infected macrophages (21) also extends to uninfected macrophages found in the spleen. Based upon the work of others showing that type II-infected macrophages secrete more IL-12 while type I/III-infected macrophages secrete less IL-12 and more IL-4 (19,21,22,60), we speculate that the strain-specific differences in secreted cytokines from infected immune cells propagates the immune cell polarization differences to uninfected immune cells (Fig. 13).

Finally, we have shown that the IIIΔ*rop16* strain is able to elicit a strong brain immune response, including infiltration of T cells and likely monocytes, despite having a very low CNS parasite burden (Fig. 6**-**8). The finding of a much stronger CNS immune response than parasite burden is consistent with prior work showing that immune cells can and do infiltrate into the CNS in the setting of a strong systemic immune response without brain infection or pathology (61). We suggest that our data add to the growing body of literature that the “immune privileged” status of the CNS is less absolute than previously thought and that we still have much to learn about what governs when and if immune cells infiltrate into the CNS.

## Materials and methods

### Ethics statement

All mouse studies and breeding were carried out in strict accordance with the Public Health Service Policy on Human Care and Use of Laboratory Animals. The protocol was approved by the University of Arizona Institutional Animal Care and Use Committee (#A-3248-01, protocol #12-391).

### Parasite maintenance

All parasite strains were maintained through serial passage in human foreskin fibroblast (gift of John Boothroyd, Stanford University, Stanford, CA) using DMEM, supplemented with 10% fetal bovine serum, 2mM glutagro, and 100 I.U/ml penicillin and 100 µg/ml streptomycin. Unless otherwise mentioned, previously described type II (Pruginaud) and type III (CEP) parasites expressing Cre recombinase and mCherry were used (42,44,45). For experiments with IIIΔ*rop16* strains, depending upon the IIIΔ*rop16* clone, we used either the previously mentioned strains or Pru*Δhpt* and CEP*Δhpt* strains in which the endogenous gene for hypoxanthine xanthine guanine phosphoribosyl transferase (HPT) (gift of John Boothroyd) has been deleted.

### Mice

Unless otherwise specified, the mice used in this study are Cre-reporter mice that only express a green fluorescent protein (GFP) after the cells have undergone Cre-mediated recombination (43). Mice were purchased from Jackson Laboratories (stock # 007906) and bred in the University of Arizona Animal Center. We used these mice in combination with our Cre-secreting parasites as a way to identify the immune cells that had been injected with parasite rhoptry proteins (42). In addition, breeding pairs of mice lacking Irgm1 and Irgm3 were provided to us courtesy of Greg Taylor (Duke University, Durham, NC) and subsequently bred in the University of Arizona Animal Center. Mice were inoculated intraperitoneally with 10,000 freshly syringe-lysed parasites diluted in 200 µl of UPS grade PBS.

### Tissue preparation for histology and protein/DNA extraction

At appropriate times post infection, mice were anesthetized with a ketamine (24 mg/ml) and xylazine (4.8 mg/ml) cocktail and transcardially perfused with ice cold phosphate buffered saline. As previously described, after harvesting organs, the left half of the mouse brain was fixed in 4% paraformaldehyde in phosphate buffer, kept at 4°C overnight, rinsed in PBS, and then was embedded in 30% sucrose (42,45). Post fixation and sucrose embedding, the brain was sectioned into 40 µm thick sagittal sections using a freezing sliding microtome (Microm HM 430). Sections were stored as free-floating sections in cryoprotective media (0.05 M sodium phosphate buffer containing 30% glycerol and 30% ethylene glycol) until stained and mounted on slides. The right half of the brain was sectioned coronally into 2 halves and stored in separate tubes. These tubes were flash frozen and stored at −80°C until used for protein or DNA extraction.

### Immunohistochemistry

As described previously, free-floating tissue sections were stained using a standard protocol (62). Brain sections were stained using the following primary antibodies: polyclonal rabbit anti-Iba-1 (019-19741, Wako Pure Chemical Industries, Ltd., (1:3000); monoclonal hamster anti-mouse CD3ε 500A2 (550277, BD Pharmingen^TM^, (1:300). Following incubation with primary antibody, sections were incubated in appropriate secondary antibodies: biotinylated goat anti-rabbit (BA-1000, Vector Laboratories (1:500) and biotinylated goat anti-hamster (BA-9100, Vector Laboratories, (1:500). Next, sections were incubated in ABC (32020, Thermo Fisher) for 1 hr followed by 3,3’-Diaminobenzidine (DAB) (SK-4100, Vector Laboratories) detection of biotinylated antibodies.

### Immune cell quantification

Brain sections stained for anti-CD3 or anti-Iba-1 antibody and detected using DAB were analyzed using light microscopy. For each brain section, twelve fields of view (FOV) throughout the cortex region of the brain beginning with the rostral region and moving caudally were sampled in a stereotyped way and as previously described (35,62). The number of CD3ε cells/FOV was quantified using SimplePCI software (Hamamatsu, Sewickley, PA) on an Olympus IMT-2 inverted light microscope (35). The number of Iba-1^+^ cells/FOV was quantified by manually counting cells with FIJI software (63). These analyses was performed on 3 brain sections per mouse, after which the resulting numbers were then averaged to obtain the average number of Iba1^+^ or CD3^+^ immune cells/brain section/mouse. Investigators quantifying CD3^+^ and Iba-1^+^ cells were blinded to the infection status of the mouse until after the data were collected.

### Protein Extraction, Quantification and Multiplex LUMINEX Assay

The caudal quarter of the flash frozen brain tissue was homogenized in radioimmunoprecipitation assay buffer (1% Triton-X, 0.1% SDS, 1X PBS) and 1% phosphatase inhibitor cocktail (P5726-1ML, Sigma-Aldrich) and 1% protease inhibitor cocktail (P8340-1ML, Sigma-Aldrich). As previously described, the samples were sonicated on ice in 3s bursts until homogenized and then were centrifuged at 4°C (62). The protein concentration of each sample was measured using Direct Detect^®^ Infrared Spectrometer. Each sample was stored at −80°C until the LUMINEX^®^ assay was performed. Cytokines and chemokines were assessed using the MILLIPLEX-MAP-Mouse-Cytokiine/Chemokine-Magnetic-Bead-Panel (MCYTOMAG-70K, EMD Millipore). This multiplex panel allows the detection of 25 different cytokines/chemokines and includes individual quality controls for each cytokine/chemokine. The samples were plated as duplicates and the plate was analyzed using a LUMINEX MAGPIX xPONENT 1.2 System which uses Milliplex Analyst software and Luminex^®^ technology to detect individual cytokine/chemokine quantities.

### Quantitative real time PCR

For quantification of parasite burden, genomic DNA from the rostral quarter of the frozen brain was isolated using DNeasy Blood and Tissue kit (69504, Qiagen) and following the manufacturer’s protocol. The *Toxoplasma* specific, highly conserved 35-repeat B1 gene was amplified using SYBR Green fluroscence detection with the Eppendorf Mastercycler ep realplex 2.2 system using primers listed in **Table S3**. GAPDH was used as house-keeping gene to normalize parasite DNA levels. Results were calculated as previously described (35,62). For quantification of Arg-1, IL-4, and IL-12, total RNA from sorted M1 and M2 cells was extracted with TRIzol™ (Life Technologies, Grand Island, NY) and according to the manufacturer’s instructions. First strand cDNA synthesis was performed using a High-Capacity cDNA Reverse Transcription kit (4368814, ThermoFisher) and following the manufacturer’s instructions. Arg-1, IL-4, and IL-12 were amplified using SYBR Green fluorescence detection with the Eppendorf Mastercycler ep realplex 2.2 system using the primers are listed in **Table S3** (21,22,41). GAPDH was used as the house-keeping gene for normalization. The reaction condition were as follows: 2 min at 50°C, 2 min at 90°C, followed by 40 cycles of 15 sec at 95°C, 15 sec at 58°C, and 1 min at 72°C, followed by a melting curve analysis.

### Cyst counts

Sagittal brain sections were washed and blocked in 3% Goat Serum in 0.3% TritonX-100/TBS for 1 hr. Sections were then incubated with biotinylated Dolichos biflorus agglutinin (DBA) (Vector Laboratories 1031, 1:500), which binds to the cyst wall (64–66). The following secondary was used: 405 Streptavidin (Invitrogen, 1:2000). Sections were mounted as previously described (35). The number of cysts were enumerated using a standard epifluorescent microscope (EVOS microscope). Only objects that expressed mCherry and stained for DBA were quantified as cysts.

### Flow Cytometry

At appropriate times post infection, mice were euthanized by CO_2_ and intracardially perfused with 20 mL ice-cold PBS, after which spleens and brains were harvested. These tissues were then made into single cell suspensions. For brains, mononuclear cells were isolated by mincing the tissue and then passing it serially through an 18-gauge needle and then a 22-gauge needle in complete RPMI (86% RPMI, 10% FBS, 1% penicillin/streptomycin, 1% L-glutamine, 1% NEAA, 1% sodium pyruvate and <0.01% β-mercaptoethanol) as described previously (67). After syringe passage, the cell suspension was passed through a 70 µm strainer and mononuclear cells were isolated using a density gradient that consisted of a 60% Percoll solution in cRPMI overlayed with a 30% Percoll solution in PBS. Brain mononuclear cells were isolated from the interphase. Spleens were made into single cell suspension and passed through a 40 µm strainer (68). Red blood cells were lysed by using ammonium chloride-potassium carbonate (ACK) lysis buffer. The total numbers of viable cells from brain and spleen suspensions were determined by trypan blue exclusion and counting on a hemocytometer. Brain and spleen samples were split in order to stain with either the T cell panel or the macrophage panel. Brain and spleen single cell suspension had F_c_ receptors blocked with 2.4G2 to prevent non-specific staining. The following directly conjugated antibodies were utilized for flow cytometry analysis of T cells: CD3 APC eFluor^®^ 780 (clone 17A2; eBioscience, 47-0032-80), CD8a PerCP-Cy 5.5 (clone 53-6.7; eBioscience, 45-0081-82), CD4 PECy7 (clone GK1.5; BioLegend, 100422), CD44 Alexa Fluor^®^ 700 (clone IM7; eBioscience, 12-0441-82), CD279 (PD-1) eFluor^®^ 450 (clone J43; eBioscience, 48-9985-82) were used to incubate cells for 30 min protected from light. The following directly conjugated antibodies were utilized for flow cytometry analysis of macrophages/microglia: CD45 PerCP-Cy5.5 (clone 30-F11; eBioscience, 45-0451), F4/80 Alexa Fluor^®^ 700 (clone BM8; BioLegend, 123130), CD11b Pacific Blue™ (clone M1/70; BioLegend, 101224), CD11c PE/Cy7 (clone N418; BioLegend, 117318), CD11c FITC (clone N418; eBioScience, 11-0114-85), CD183 (CXCR3) phycoerythrin (PE) (clone CXCR3-173; BioLegend, 126505), CD183 (CXCR3) PE/Cy7 (clone CXCR3-173; BioLegend, 126516), CD206 (MMR) phycoerythrin (PE) (clone C068C2; BioLegend, 141706), CD80 APC (clone 16-10A1; eBioscience 17-0801-82), CD80 PE/Cy5 (clone 16-10A1; BioLegend, 104712), CD86 APC (clone GL-1; BioLegend, 105012), CD86 PE/Cy5 (clone GL-1; BioLegend, 105016), Ly-6G/Ly-6C (Gr-1) PE/Cy5 (clone RB6-8C5; BioLegend, 108410). Cells were incubated with appropriate antibodies for 30 min, while being protected from light. After surface staining, cells were then stained with live/dead Fixable Yellow Dead Cell Stain Kit (Life Technologies, L34959) for 30 min to distinguish between live and dead cells. For intracellular cytokine staining, while protected from light, samples were washed, permeabilized, and fixed using a permeabilization and fixation kit (eBioscience, 00-5223-56; 00-5123-43; 00-8333-56). An intracellular staining protocol was used to stain for FoxP3 PE (clone FJK-16s; eBioscience, 12-5773-82), and IFN-γ APC (clone XMG1.2; eBioscience 17-7311-82) for 30 min. Samples were washed after each staining step to remove residual unbound antibody. A BD LSR II (BD Biosciences, San Jose, CA; University of Arizona Cancer Center) was used to run the samples and Flowjo (Treestar) was used for all flow cytometry analysis.

For sorting, splenocytes were isolated as described above and stained with the macrophage panel followed by the live/dead staining as denoted above. Samples were washed and resuspended in PBS until sorted, which was done on the same day as isolation. A FACS Aria III (BD Biosciences, San Jose, CA; University of Arizona Cancer Center) was used to sort M1 and M2 populations. After sorting, samples were resuspended in TRIzol™ (Life Technologies, Grand Island, NY) and stored at −80°C until RNA extraction.

### Plasmid construction

All the plasmids and primers used to make and validate the IIIΔ*rop16* strain are listed in **Table S3**. The *rop16*-targeting CRISPR plasmids (sg *rop16*Up and sg *rop16*Down) were constructed from sgUPRT using a previously described Q5 mutagenesis protocol (51–53). To generate a plasmid for inserting *hxgprt* into the *rop16* locus, upstream (500-bp) and downstream (500-bp) regions directly adjacent to the sgROP16Up and sg *rop16*Down target sequence were used to flank *hxgprt* via sequential restriction cloning.

### Generation of IIIΔ*rop16* knockout and IIIΔ*rop16::ROP16III*

To disrupt *rop16* in type IIIΔ*hpt*, we transfected parasites with 3 plasmids: the sg *rop16*Up CRISPR and sg *rop16*Down CRISPR plasmids and the pTKO (69) plasmid containing *rop16* homology regions surrounding a selectable marker (*hxgprt*) or with the toxofilin-Cre cassette (44) (**Fig. S5**). Selection by growth for 4 to 8 days in 25 mg/ml mycophenolic acid and 50 mg/ml xanthine (70) was used to obtain stably resistant clones with *hxgprt* integration. These clones were subsequently screened by PCR to confirm disruption of the *rop16* locus (**Fig. S5**). Clones negative for *rop16* and positive for integration of *hxgprt* were confirmed by western blot to have lost the *rop16*-dependent phosphorylation of STAT6 (59). In addition, clones with the toxofilin-Cre cassette were confirmed to trigger Cre-mediated recombination as previously described (44).

To complement *rop16*, IIIΔ*rop16* parasites were transfected with 50 μgs of linearized plasmid DNA harboring a FLAG-tagged *ROP16_III_* gene and a bleomycin resistance marker. Post transfection freshly egressed parasites were resuspended in DMEM supplemented with 50 μg/mL of Zeocin (InvivoGEN, 11006-33-0) for 4 hour and then added to HFF monolayers supplemented with 5μg/mL Zeocin to select for integrants. This process was repeated 3 times prior to plating by limiting dilution to isolate single clones. Single clones were subsequently screened by PCR for *rop16* integration (**Fig. S5**). The IIIΔ*rop16::ROP16_III_* clones were all derived from the IIIΔ*rop16* strain that expresses toxofilin-Cre.

### Peritoneal Exudate Cells Isolation

Cre-reporter mice or Irgm1/3 knockout mice were infected with type II, type III, IIIΔ*rop16*, or IIIΔ*rop16::ROP16_III_*. At appropriate times, peritoneal exudate cells (PECs) were collected by peritoneal lavage with 10 ml of cold 1 x PBS. PECs were incubated in FcBlock for 10 min as described above. PECs were subsequently stained with CD45 marker, followed by with live/dead staining as described above, and then analyzed using a BD LSR II (BD Biosciences, San Jose, CA; University of Arizona Cancer Center) (42).

### Statistical analysis

Statistical analyses were performed using Prism 7.0 software. To improve distributional characteristics, total numbers of CD3 and Iba-1 cells were log transformed prior to analysis. Unless otherwise specified, two-way analysis of variance (ANOVA) with Fisher’s protected LSD was used, with the cohort as the block factor and parasite strain as the experimental factor. For cytokine levels, the data were analyzed using a one-way ANOVA with Bonferroni’s post-hoc test.

## Acknowledgements

The authors would like to thank all members of the Koshy Lab for helpful discussions and critical review of the manuscript. We would like to thank Dean Billheimer (Director of BIO5 Statistical Consulting Group) for his suggestions for data analyses. We would also like to thank the Doyle Lab, especially Vivian Nguyen for the use of their Luminex® MAGPIX for the cytokine assays. Finally, we would like to thank Greg Taylor (Duke University), for providing us with breeding pairs for the *Irgm1/3* knockout mice. Research in this manuscript was directly supported by the UA Flow Cytometry Shared Resource funded by the National Cancer Institute (P30CA023074).

---

**Table S1. List of cytokines and chemokines from the 25-plex LUMINEX assay.** The table shows the mean concentration (pg/ml) ± SEM of cytokines and chemokines. Blue represents those cytokines or chemokines with a ≥2-fold change over saline treated controls. p-values are based on one-way ANOVA with Bonferroni post-hoc test.

**Fig S1. Gating scheme for macrophage markers.** Immune cells were isolated from the brain and stained for macrophage markers. Single cells were discriminated from doublets by plotting side scatter height (SSC-H) versus side scatter area (SSC-A). Cells were selected by plotting SSC-A versus forward scatter area (FSC-A). Live cells were gated on live/dead Yellow^-^. CD45^+^ CD3^-^ cells were gated by plotting CD3 versus CD45. From the CD45^+^ gate, F4/80^+^ and F4/80^-^ cells were gated by plotting FSC-A versus F4/80. From the F4/80^+^ gate, macrophages (Macs) were gated by plotting CD11c versus CD11b. From the Macs gate, (CD80^+^/CD86^+^) M1 cells were gated by plotting CD80/CD86 versus CD11b. From the Macs gate, (MMR^+^/CxCR3^+^) M2 cells were gated by plotting MMR/CXCR3 versus CD11b. Uninfected controls and isotype controls were used to establish the gating scheme.

**Fig S2. Gating scheme for T cell markers.** Immune cells were isolated from the brain and stained for T cell markers. Single cells were discriminated from doublets by plotting side scatter height (SSC-H) versus side scatter area (SSC-A). Cells were selected by plotting SSC-A versus forward scatter area (FSC-A). Live cells were gated on live/dead Yellow^-^. CD3^+^ cells were gated by plotting SSC-A versus CD3. From the CD3^+^ gate, CD4^+^ and CD8^+^ cells were gated by plotting CD4 versus CD8. From the CD4^+^ gate, FoxP3^+^ Tregs were gated by plotting FoxP3 versus CD4. Uninfected controls and isotype controls were used to establish the gating scheme.

**Fig S3. Placing CD80/CD86 or MMR/CXCR3 in the same or individual channels results in similar findings in type II or type III-infected mice.** At 21 dpi, immune cells were isolated from the CNS of either type II or type III-infected mice, split, stained for macrophage markers, and then analyzed by flow cytometry. **A.** For type II-infected mice, the percentage and number of M2 macrophages identified by placing MMR and CXCR3 in the same channel or separate channels. **B.** For type II infected mice, the percentage and number of M1 macrophages identified by placing CD80 and CD86 in the same channel or separate channels. **C.** As in (**A**) except for type III-infected mice. **D.** As in (**B**) except for type III-infected mice. Bars, mean ± SEM. N= 5 mice/infected group. ns = not significant, non-parametric t-test.

**Figure S4. IL-12 expression and IFN-γ production are higher in M1s whereas Arg-1 and IL-4 expression is higher in M2s.** Mice were inoculated with type II or type III parasites. At 5dpi, splenocytes were isolated, stained, and sorted into M1s and M2s. Q-PCR was performed on RNA isolated from the M1s and M2s. **A,B.** Q-PCR quantification of IL-12, Arg-1, and IL-4 expression from M1s and M2s from type II-infected mice. **C, D.** As in (A,B) except from M1s and M2s from type III-infected mice. **E.** Quantification of the frequency of IFN-γ producing M1s and M2s. **F.** Quantification of the mean fluorescent intensity of IFN-γ in M1s and M2s. N =5 Mice/group

**Table S2: List of cells types characterized between type II and type III-infected mice.**

**Figure S5. Generation and confirmation of IIIΔ*rop16* and IIIΔ*rop16*::*ROP16_III_*. A.** Schematic representation of the approach used to create the IIIΔ*rop16* and IIIΔ*rop16::ROP16_III_* complemented strains. Type IIIΔ*hpt* parasites were transfected with CRISPR/CAS9 vectors targeting 500bp upstream (gRNA Up) and downstream (gRNA Down) of the *rop*16 coding sequence and a linearized vector with 500bp regions of homology (HR) to the 5’ and 3’UTRs of *rop16* surrounding either the selectable marked *HXGPRT* alone (not shown) or the selectable marked *HXGPRT* and the *toxofilin-Cre* coding sequence (shown). Complementation was achieved using a linearized vector encoding a FLAG-tagged *ROP16* and a selectable bleomycin-resistance marker. **B.** PCR of the entire *rop16* locus for the IIIΔ*rop16* and IIIΔ*rop16::ROP16_III_* strains. PCR analysis of SAG1 was used as a DNA control. **C.** Western blots from HFFs stimulated with IL-4 or infected with parental (Type III), IIIΔ*rop16*, or IIIΔ*rop16::ROP16_III_* parasites. Protein isolation was done at 18 hours post-infection or stimulation. HFFs were infected at a MOI of 5.

**Table S3. List of Primers used throughout the paper.**

## REFERENCES

1. Suzuki Y. Immunopathogenesis of Cerebral Toxoplasmosis. J Infect Dis. 2002 Dec 1;186:S234–40.

2. Carruthers VB. Host cell invasion by the opportunistic pathogen Toxoplasma gondii. Acta Trop. 2002 Feb;81(2):111–22.

3. Hill D, Dubey JP. Toxoplasma gondii: transmission, diagnosis and prevention. Clin Microbiol Infect Off Publ Eur Soc Clin Microbiol Infect Dis. 2002 Oct;8(10):634–40.

4. Wreghitt TG, Gray JJ, Pavel P, Balfour A, Fabbri A, Sharples LD, et al. Efficacy of pyrimethamine for the prevention of donor-acquired Toxoplasma gondii infection in heart and heart-lung transplant patients. Transpl Int Off J Eur Soc Organ Transplant. 1992 Sep;5(4):197–200.

5. Ruskin J, Remington JS. Toxoplasmosis in the compromised host. Ann Intern Med. 1976 Feb;84(2):193–9.

6. Luft BJ, Remington JS. Toxoplasmic encephalitis in AIDS. Clin Infect Dis Off Publ Infect Dis Soc Am. 1992 Aug;15(2):211–22.

7. Desmonts G, Couvreur J. Congenital toxoplasmosis. A prospective study of 378 pregnancies. N Engl J Med. 1974 May 16;290(20):1110–6.

8. Grigg ME, Ganatra J, Boothroyd JC, Margolis TP. Unusual abundance of atypical strains associated with human ocular toxoplasmosis. J Infect Dis. 2001 Sep 1;184(5):633–9.

9. Kamerkar S, Davis PH. Toxoplasma on the Brain: Understanding Host-Pathogen Interactions in Chronic CNS Infection. J Parasitol Res [Internet]. 2012 [cited 2013 Aug 10];2012. Available from: http://www.ncbi.nlm.nih.gov/pmc/articles/PMC3321570/

10. Ferreira IMR, Vidal JE, de Mattos C de CB, de Mattos LC, Qu D, Su C, et al. Toxoplasma gondii isolates: multilocus RFLP-PCR genotyping from human patients in Sao Paulo State, Brazil identified distinct genotypes. Exp Parasitol. 2011 Oct;129(2):190–5.

11. Dubey JP, Lago EG, Gennari SM, Su C, Jones JL. Toxoplasmosis in humans and animals in Brazil: high prevalence, high burden of disease, and epidemiology. Parasitology. 2012 Sep;139(11):1375–424.

12. Furtado JM, Winthrop KL, Butler NJ, Smith JR. Ocular toxoplasmosis I: parasitology, epidemiology and public health. Clin Experiment Ophthalmol. 2013 Feb;41(1):82–94.

13. Remington JS, McLeod R, Wilson CB, Desmonts G. CHAPTER 31 - Toxoplasmosis. In: Infectious Diseases of the Fetus and Newborn (Seventh Edition) [Internet]. Philadelphia: W.B. Saunders; 2011. p. 918–1041. Available from: https://www.sciencedirect.com/science/article/pii/B9781416064008000316

14. Boothroyd JC, Grigg ME. Population biology of Toxoplasma gondii and its relevance to human infection: do different strains cause different disease? Curr Opin Microbiol. 2002 Aug 1;5(4):438–42.

15. Sibley LD, Boothroyd JC. Virulent strains of Toxoplasma gondii comprise a single clonal lineage. Nature. 1992 Sep 3;359(6390):82–5.

16. Howe DK, Sibley LD. Toxoplasma gondii comprises three clonal lineages: correlation of parasite genotype with human disease. J Infect Dis. 1995 Dec;172(6):1561–6.

17. Su C, Khan A, Zhou P, Majumdar D, Ajzenberg D, Dardé M-L, et al. Globally diverse Toxoplasma gondii isolates comprise six major clades originating from a small number of distinct ancestral lineages. Proc Natl Acad Sci. 2012 Apr 10;109(15):5844–9.

18. Saeij JPJ, Boyle JP, Coller S, Taylor S, Sibley LD, Brooke-Powell ET, et al. Polymorphic secreted kinases are key virulence factors in toxoplasmosis. Science. 2006 Dec 15;314(5806):1780–3.

19. Butcher BA, Fox BA, Rommereim LM, Kim SG, Maurer KJ, Yarovinsky F, et al. Toxoplasma gondii rhoptry kinase ROP16 activates STAT3 and STAT6 resulting in cytokine inhibition and arginase-1-dependent growth control. PLoS Pathog. 2011 Sep;7(9):e1002236.

20. Rosowski EE, Lu D, Julien L, Rodda L, Gaiser RA, Jensen KDC, et al. Strain-specific activation of the NF-kappaB pathway by GRA15, a novel Toxoplasma gondii dense granule protein. J Exp Med. 2011 Jan 17;208(1):195–212.

21. Jensen KDC, Wang Y, Wojno EDT, Shastri AJ, Hu K, Cornel L, et al. Toxoplasma polymorphic effectors determine macrophage polarization and intestinal inflammation. Cell Host Microbe. 2011 Jun 16;9(6):472–83.

22. Robben PM, Mordue DG, Truscott SM, Takeda K, Akira S, Sibley LD. Production of IL-12 by macrophages infected with Toxoplasma gondii depends on the parasite genotype. J Immunol Baltim Md 1950. 2004 Mar 15;172(6):3686–94.

23. Suzuki Y, Conley FK, Remington JS. Differences in virulence and development of encephalitis during chronic infection vary with the strain of Toxoplasma gondii. J Infect Dis. 1989 Apr;159(4):790–4.

24. Suzuki Y, Joh K. Effect of the strain of Toxoplasma gondii on the development of toxoplasmic encephalitis in mice treated with antibody to interferon-gamma. Parasitol Res. 1994;80(2):125–30.

25. Fentress SJ, Behnke MS, Dunay IR, Mashayekhi M, Rommereim LM, Fox BA, et al. Phosphorylation of immunity-related GTPases by a Toxoplasma gondii-secreted kinase promotes macrophage survival and virulence. Cell Host Microbe. 2010 Dec 16;8(6):484–95.

26. Niedelman W, Gold DA, Rosowski EE, Sprokholt JK, Lim D, Arenas AF, et al. The Rhoptry Proteins ROP18 and ROP5 Mediate Toxoplasma gondii Evasion of the Murine, But Not the Human, Interferon-Gamma Response. PLOS Pathog. 2012 Jun 28;8(6):e1002784.

27. Henry SC, Daniell XG, Burroughs AR, Indaram M, Howell DN, Coers J, et al. Balance of Irgm protein activities determines IFN-γ-induced host defense. J Leukoc Biol. 2009;85(5):877–85.

28. Suzuki Y, Orellana MA, Schreiber RD, Remington JS. Interferon-gamma: the major mediator of resistance against Toxoplasma gondii. Science. 1988 Apr 22;240(4851):516–8.

29. Suzuki Y, Conley FK, Remington JS. Treatment of toxoplasmic encephalitis in mice with recombinant gamma interferon. Infect Immun. 1990 Sep 1;58(9):3050–5.

30. Suzuki Y, Claflin J, Wang X, Lengi A, Kikuchi T. Microglia and macrophages as innate producers of interferon-gamma in the brain following infection with Toxoplasma gondii. Int J Parasitol. 2005 Jan;35(1):83–90.

31. Sa Q, Ochiai E, Tiwari A, Perkins S, Mullins J, Gehman M, et al. Cutting Edge: IFN-γ Produced by Brain-Resident Cells Is Crucial To Control Cerebral Infection with Toxoplasma gondii. J Immunol Baltim Md 1950. 2015 Aug 1;195(3):796–800.

32. Burg JL, Grover CM, Pouletty P, Boothroyd JC. Direct and sensitive detection of a pathogenic protozoan, Toxoplasma gondii, by polymerase chain reaction. J Clin Microbiol. 1989 Aug;27(8):1787–92.

33. Buchbinder S, Blatz R, Christian Rodloff A. Comparison of real-time PCR detection methods for B1 and P30 genes of Toxoplasma gondii. Diagn Microbiol Infect Dis. 2003 Apr;45(4):269–71.

34. Noor S, Habashy AS, Nance JP, Clark RT, Nemati K, Carson MJ, et al. CCR7-dependent immunity during acute Toxoplasma gondii infection. Infect Immun. 2010 May;78(5):2257–63.

35. Cekanaviciute E, Dietrich HK, Axtell RC, Williams AM, Egusquiza R, Wai KM, et al. Astrocytic TGF-β signaling limits inflammation and reduces neuronal damage during central nervous system Toxoplasma infection. J Immunol Baltim Md 1950. 2014 Jul 1;193(1):139–49.

36. Knoll LJ, Boothroyd JC. Isolation of developmentally regulated genes from Toxoplasma gondii by a gene trap with the positive and negative selectable marker hypoxanthine-xanthine-guanine phosphoribosyltransferase. Mol Cell Biol. 1998 Feb;18(2):807–14.

37. O’Brien CA, Overall C, Konradt C, O’Hara Hall AC, Hayes NW, Wagage S, et al. CD11c-Expressing Cells Affect Regulatory T Cell Behavior in the Meninges during Central Nervous System Infection. J Immunol Baltim Md 1950. 2017 15;198(10):4054–61.

38. Hwang YS, Shin J-H, Yang J-P, Jung B-K, Lee SH, Shin E-H. Characteristics of Infection Immunity Regulated by Toxoplasma gondii to Maintain Chronic Infection in the Brain. Front Immunol. 2018;9:158.

39. Murray PJ, Allen JE, Biswas SK, Fisher EA, Gilroy DW, Goerdt S, et al. Macrophage activation and polarization: nomenclature and experimental guidelines. Immunity. 2014 Jul 17;41(1):14–20.

40. Jin RM, Blair SJ, Warunek J, Heffner RR, Blader IJ, Wohlfert EA. Regulatory T Cells Promote Myositis and Muscle Damage in Toxoplasma gondii Infection. J Immunol Baltim Md 1950. 2017 01;198(1):352–62.

41. Nance JP, Vannella KM, Worth D, David C, Carter D, Noor S, et al. Chitinase dependent control of protozoan cyst burden in the brain. PLoS Pathog. 2012;8(11):e1002990.

42. Koshy AA, Dietrich HK, Christian DA, Melehani JH, Shastri AJ, Hunter CA, et al. Toxoplasma Co-opts Host Cells It Does Not Invade. PLoS Pathog. 2012 Jul 26;8(7):e1002825.

43. Madisen L, Zwingman TA, Sunkin SM, Oh SW, Zariwala HA, Gu H, et al. A robust and high-throughput Cre reporting and characterization system for the whole mouse brain. Nat Neurosci. 2010 Jan;13(1):133–40.

44. Koshy AA, Fouts AE, Lodoen MB, Alkan O, Blau HM, Boothroyd JC. Toxoplasma secreting Cre recombinase for analysis of host-parasite interactions. Nat Methods. 2010 Apr;7(4):307–9.

45. Cabral CM, Tuladhar S, Dietrich HK, Nguyen E, MacDonald WR, Trivedi T, et al. Neurons are the Primary Target Cell for the Brain-Tropic Intracellular Parasite Toxoplasma gondii. PLoS Pathog. 2016 Feb;12(2):e1005447.

46. Jensen KDC, Hu K, Whitmarsh RJ, Hassan MA, Julien L, Lu D, et al. Toxoplasma gondii Rhoptry 16 Kinase Promotes Host Resistance to Oral Infection and Intestinal Inflammation Only in the Context of the Dense Granule Protein GRA15. Infect Immun. 2013 Jun 1;81(6):2156–67.

47. Xu S, Shinohara ML. Tissue-Resident Macrophages in Fungal Infections. Front Immunol. 2017;8:1798.

48. Roberts CA, Dickinson AK, Taams LS. The Interplay Between Monocytes/Macrophages and CD4+ T Cell Subsets in Rheumatoid Arthritis. Front Immunol [Internet]. 2015 [cited 2019 Mar 3];6. Available from: https://www.frontiersin.org/articles/10.3389/fimmu.2015.00571/full

49. Mojica FJM, Díez-Villaseñor C, García-Martínez J, Soria E. Intervening Sequences of Regularly Spaced Prokaryotic Repeats Derive from Foreign Genetic Elements. J Mol Evol. 2005 Feb 1;60(2):174–82.

50. Ran FA, Hsu PD, Wright J, Agarwala V, Scott DA, Zhang F. Genome engineering using the CRISPR-Cas9 system. Nat Protoc Lond. 2013 Nov;8(11):2281–308.

51. Shen B, Brown KM, Lee TD, Sibley LD. Efficient gene disruption in diverse strains of Toxoplasma gondii using CRISPR/CAS9. mBio. 2014 May 13;5(3):e01114–01114.

52. Sidik SM, Hackett CG, Tran F, Westwood NJ, Lourido S. Efficient Genome Engineering of Toxoplasma gondii Using CRISPR/Cas9. PLoS One San Franc. 2014 Jun;9(6):e100450.

53. Shen B, Brown K, Long S, Sibley LD. Development of CRISPR/Cas9 for Efficient Genome Editing in Toxoplasma gondii. Methods Mol Biol Clifton NJ. 2017;1498:79–103.

54. Zhao YO, Khaminets A, Hunn JP, Howard JC. Disruption of the Toxoplasma gondii Parasitophorous Vacuole by IFNγ-Inducible Immunity-Related GTPases (IRG Proteins) Triggers Necrotic Cell Death. PLoS Pathog. 2009 Feb 6;5(2):e1000288.

55. Haldar AK, Saka HA, Piro AS, Dunn JD, Henry SC, Taylor GA, et al. IRG and GBP Host Resistance Factors Target Aberrant, “Non-self” Vacuoles Characterized by the Missing of “Self” IRGM Proteins. PLOS Pathog. 2013 Jun 13;9(6):e1003414.

56. Coers J, Gondek DC, Olive AJ, Rohlfing A, Taylor GA, Starnbach MN. Compensatory T Cell Responses in IRG-Deficient Mice Prevent Sustained Chlamydia trachomatis Infections. PLOS Pathog. 2011 Jun 23;7(6):e1001346.

57. Coers J, Bernstein-Hanley I, Grotsky D, Parvanova I, Howard JC, Taylor GA, et al. Chlamydia muridarum Evades Growth Restriction by the IFN-γ-Inducible Host Resistance Factor Irgb10. J Immunol. 2008 May 1;180(9):6237–45.

58. Feng CG, Zheng L, Jankovic D, Báfica A, Cannons JL, Watford WT, et al. The immunity-related GTPase Irgm1 promotes the expansion of activated CD4^+^ T cell populations by preventing interferon-γ-induced cell death. Nat Immunol. 2008 Nov;9(11):1279–87.

59. Saeij JPJ, Coller S, Boyle JP, Jerome ME, White MW, Boothroyd JC. Toxoplasma co-opts host gene expression by injection of a polymorphic kinase homologue. Nature. 2007 Jan 18;445(7125):324–7.

60. Hunter CA, Sibley LD. Modulation of innate immunity by Toxoplasma gondii virulence effectors. Nat Rev Microbiol. 2012 Nov;10(11):766–78.

61. Hickey WF, Kimura H. Graft-vs.-host disease elicits expression of class I and class II histocompatibility antigens and the presence of scattered T lymphocytes in rat central nervous system. Proc Natl Acad Sci. 1987 Apr 1;84(7):2082–6.

62. Cabral CM, McGovern KE, MacDonald WR, Franco J, Koshy AA. Dissecting Amyloid Beta Deposition Using Distinct Strains of the Neurotropic Parasite Toxoplasma gondii as a Novel Tool. ASN Neuro. 2017 Aug;9(4):1759091417724915.

63. Schindelin J, Arganda-Carreras I, Frise E, Kaynig V, Longair M, Pietzsch T, et al. Fiji: an open-source platform for biological-image analysis. Nat Methods. 2012 Jun 28;9(7):676–82.

64. Knoll LJ, Boothroyd JC. Isolation of Developmentally Regulated Genes from Toxoplasma gondii by a Gene Trap with the Positive and Negative Selectable Marker Hypoxanthine-Xanthine-Guanine Phosphoribosyltransferase. Mol Cell Biol. 1998 Feb;18(2):807–14.

65. Fux B, Nawas J, Khan A, Gill DB, Su C, Sibley LD. Toxoplasma gondii Strains Defective in Oral Transmission Are Also Defective in Developmental Stage Differentiation. Infect Immun. 2007 May;75(5):2580–90.

66. Schaeffer M, Han S-J, Chtanova T, Dooren GG van, Herzmark P, Chen Y, et al. Dynamic Imaging of T Cell-Parasite Interactions in the Brains of Mice Chronically Infected with Toxoplasma gondii. J Immunol. 2009 May 15;182(10):6379–93.

67. Landrith TA, Sureshchandra S, Rivera A, Jang JC, Rais M, Nair MG, et al. CD103+ CD8 T Cells in the Toxoplasma-Infected Brain Exhibit a Tissue-Resident Memory Transcriptional Profile. Front Immunol. 2017;8:335.

68. Barrigan LM, Tuladhar S, Brunton JC, Woolard MD, Chen C, Saini D, et al. Infection with Francisella tularensis LVS clpB leads to an altered yet protective immune response. Infect Immun. 2013 Jun;81(6):2028–42.

69. Pernas L, Boothroyd JC. Association of host mitochondria with the parasitophorous vacuole during Toxoplasma infection is not dependent on rhoptry proteins ROP2/8. Int J Parasitol. 2010 Oct;40(12):1367–71.

70. Donald RG, Roos DS. Insertional mutagenesis and marker rescue in a protozoan parasite: cloning of the uracil phosphoribosyltransferase locus from Toxoplasma gondii. Proc Natl Acad Sci. 1995 Jun 6;92(12):5749–53.

